# Arachidonic acid availability controls neutrophil swarm initiation and scaling

**DOI:** 10.64898/2026.04.15.718800

**Authors:** Evelyn D. Strickland, Rosheni Kandaswamy, Patrick J. Zager, Henry de Belly, Tasha K. Phillips, Alex Hopke, Orion D. Weiner

## Abstract

Neutrophils are first responders of the vertebrate immune system. To efficiently converge on sites of injury and infection, neutrophils engage in a collective migration process known as swarming, in which a small number of activated cells generate amplified recruitment of hundreds to thousands of additional neutrophils. How neutrophils initiate, scale, and terminate these swarms is not well understood. Here we define key roles for the mechanosensitive phospholipase cPLA_2_ and its product, arachidonic acid (AA), in swarm initiation and scaling. We observe that swarm-initiating neutrophils satisfy the conditions for cPLA_2_ activation through two coincident inputs: yeast-contact-mediated Ca^2+^ influx and nuclear stretch following cell spreading along fungal hyphae and clusters. This co-requirement for both chemical and physical features of pathogens may explain how neutrophils restrict swarming to insults that require collective action. We further demonstrate that AA release is necessary and sufficient for swarming and that AA levels regulate swarm magnitude. We propose that neutrophils share AA across multiple yeast-engaged cells to collectively assess infection magnitude. Because calcium influx, nuclear deformation, and cPLA_2_-mediated AA generation are also features of sterile-injury inflammatory responses, our findings suggest a unifying circuit for swarm regulation across injury and infection contexts.

## Introduction

Neutrophils are the innate immune system’s first responders to sites of infection and tissue injury. Initial inflammation cues are limited in range and often cannot directly recruit sufficient neutrophils for pathogen sequestration and tissue repair. To overcome this limitation, neutrophils amplify these primary signals through a collectively coordinated migration process known as swarming. Like bird flocks or ant colonies[1–3], neutrophil swarming emerges from local interactions that give rise to coordinated, large-scale movement. Neutrophil swarming is predominantly driven by the sensing and production of LTB_4_[4–6] through a highly controlled, auto-amplifying circuit[7, 8]. First observed in Toxoplasma gondii-infected lymph tissues[9], swarming has since been reported across multiple species and in diverse contexts, including intravital mouse infection/sterile injury[4, 9–16], zebrafish sterile injury/infection[6, 17, 18], mouse/human *ex vivo* infection[5, 8, 19–23], and human tissue samples[24, 25].

Swarming is not an all-or-none response but scales strongly with insult size[5, 22], and the molecular mechanisms underlying this scaling are not known. The rapid, self-amplified nature of neutrophil swarming creates a key challenge: neutrophils must rapidly scale their response to contain expanding infections and clear damaged tissue while also limiting collateral damage from their influx and release of cytotoxic agents[26–28]. How neutrophils initiate, regulate, and ultimately terminate these self-amplifying chemotactic gradients remains poorly understood. This represents a fundamental gap in our understanding of how collective immune responses are controlled.

Arachidonic acid (AA) is the limiting precursor for the production of the key swarming signal LTB_4_. In neutrophils, LTB_4_ exposure alone fails to elicit significant LTB_4_ generation[29–31] unless free AA is available, establishing AA as a required co-input for the LTB_4_ positive feedback circuit[31–34]. During tissue injury, AA is generated by epithelial cells as injury-induced nuclear swelling and calcium influx activate the mechanosensitive cytosolic phospholipase A2 (cPLA_2_)[35–38]. Because this same enzyme is required for neutrophil LTB_4_ generation[39–42], we hypothesized that neutrophils similarly integrate nuclear deformation and calcium influx to control their own AA release for swarm initiation and scaling.

In contrast to injury-mediated arachidonic acid production, where hypotonic swelling of epithelial nuclei controls cPLA_2_ activation, we hypothesized that distinct mechanical triggers govern cPLA_2_ activation for infection-based swarms. Specifically, the nuclear stretch required for cPLA_2_ activation could arise by the engagement of pathogens that are too large for a single neutrophil to phagocytose[23, 43]. Consistent with this idea, interactions with oversized targets such as hyphae or spherical yeast clusters trigger robust swarming, whereas individual spherical yeast (blastospores) do not[5, 8, 22, 23, 44]. Here we analyze human primary neutrophils in a simplified *ex vivo* context to define how nuclear deformation contributes to swarm initiation. In parallel, we manipulate the levels and spatial dynamics of arachidonic acid to demonstrate its central role in swarm initiation and scaling. Our work suggests that mechanically triggered cPLA_2_ activity is a unifying regulator of swarm initiation and scaling in both injury and infection contexts.

## Results

### Neutrophil swarms scale with infection material cluster size

If physical stretch is a key determinant for swarm initiation, we would expect larger amounts of infectious material (i.e. more surface area for swarm-initiating cells to stretch across) to elicit stronger swarming responses **(Figure 1A)**. In previous work, we leveraged a reductionist assay to study infection-mediated neutrophil swarms and analyze swarming wave propagation at the millimeter scale[8, 23]. Using micro-arrayed dots of a highly charged polymer on imaging glass, we created large arrays of roughly circular ‘target’ clusters of heat-killed *Candida Albicans* **(Figure 1 – Supplement 1A)**[45]. These clusters generally span 100-200 µm in diameter and have a range of yeast densities because of natural variation in the manufacturing process **(Figure 1 – Supplement 1B, C)**. Using this assay, we found that neutrophils accumulate at targets in proportion to the number of yeast present **(Figure 1B, C)**. Our measurements are in rough agreement with previous work[5, 23] and indicate that swarming scales with the number of yeast at the site (or more likely, the density/cluster size of yeast in the area). These data indicate that neutrophil swarms scale with target size, linking stimulus magnitude to collective recruitment.

**Figure 1.**
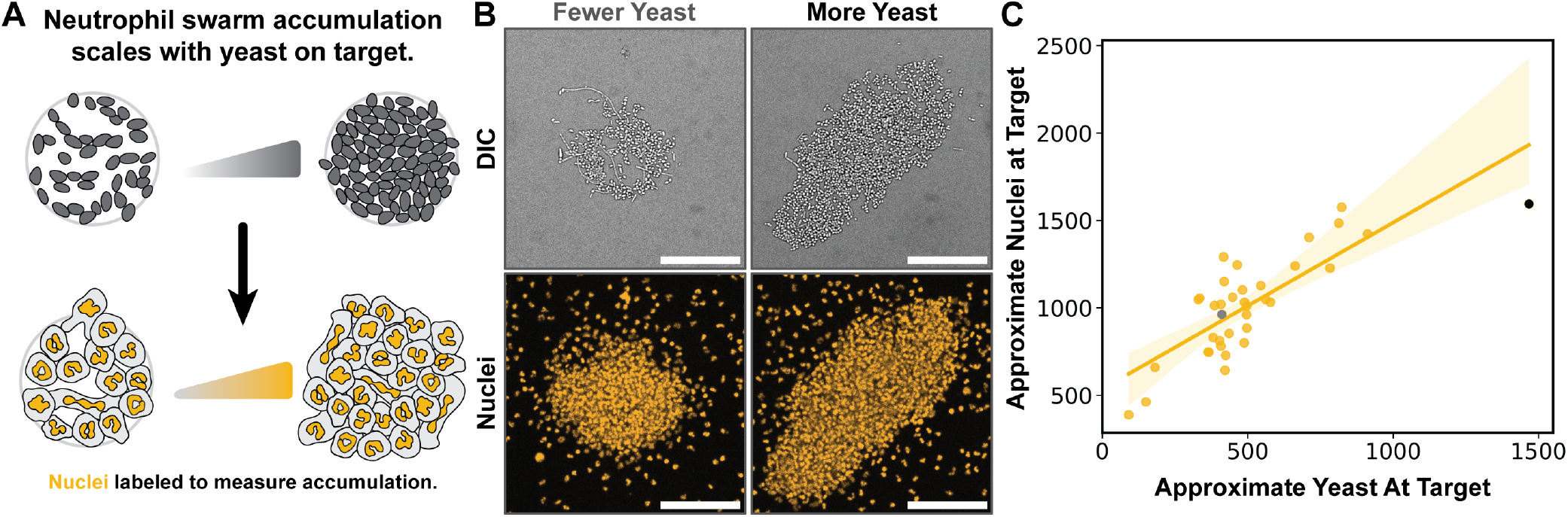
Swarm magnitude scales with target size. (**A**) Neutrophil nuclei are labeled with Hoechst 33342 to assay neutrophil accumulation at patterned mock infection sites over a range of yeast densities. (**B**) Example targets with variable yeast densities are shown in DIC before cell addition (top) with a maximum intensity projection of neutrophil nuclei that accumulate at each after 60 min (bottom, Scale bar: 100 µm; grey and black dots in **C** correspond to fewer and greater numbers of yeast at the target, respectively). (**C**) Neutrophil accumulation at the target scales proportionally with yeast number (Target n = 36, Volunteer N = 3).

### Neutrophil stretching on yeast clusters triggers swarm waves

We hypothesized that neutrophil stretching on yeast targets generates the calcium influx and nuclear mechanical deformation needed for cPLA_2_ activation and AA generation. To investigate whether swarm initiation consistently coincides with neutrophil stretch in yeast-mediated swarms, we reexamined this assay at higher magnification using calcium dyes as a live activation readout.

Following neutrophil introduction into the assay, most cells remain quiescent, but a small number touch down and patrol randomly until encountering a yeast target site. Upon target contact, neutrophils exhibit strong calcium flux as they spread across small hyphae or dense yeast clusters in a process resembling frustrated phagocytosis **(Figure 2A)**. This contact-mediated calcium influx represents one of the two co-inputs to cPLA_2_ activation. Before the explosive recruitment of hundreds of cells to the target, a series of stereotyped phases can be seen in the movies or visualized in 2D kymographs **(Figure 2B, Movie 1, Figure 2 – Supplement 1A**,**B)**. First, a patrolling neutrophil (or in some cases multiple simultaneously arriving neutrophils) contacts a yeast, exhibits a bright calcium transient, and begins to spread across multiple yeast. As one or more neutrophils stretch on the yeast target, adjacent neutrophils exhibit rapid flickering calcium transients, followed by weak local recruitment of cells to the target site. This flickering is consistent with passively diffusing signals from cells at the target to their neighbors **(Figure 2 – Supplement 1C)**. During the next phase, a “swarm-initiating” neutrophil or cluster of neutrophils generate a much more intense calcium influx; triggering a rapidly propagating, long-range series of calcium influxes in neighboring cells and resulting in the activation and recruitment of hundreds of neutrophils. We have previously established that this rapid activity wave is consistent with a self-generated LTB_4_ relay, in which activated cells trigger their neighbors to generate a rapid, self-amplifying wave of chemoattractant [8]. Such swarming waves consistently emerge closer to the average centroid of “first contact” stretching neutrophil(s), as opposed to the centroid of the target itself **(Figure 2 – Supplement 1D)**.

**Figure 2.**
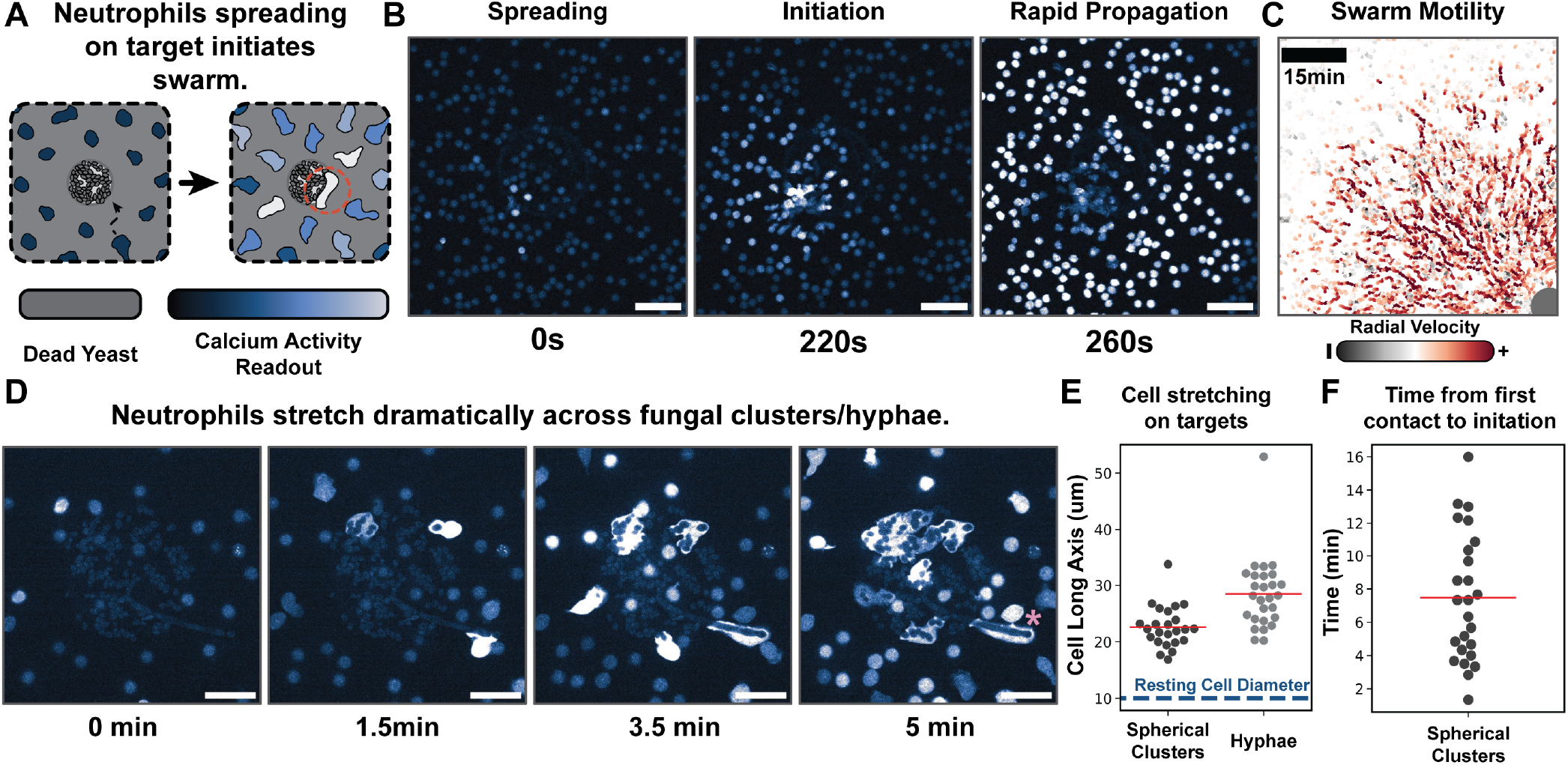
Neutrophil stretching on yeast clusters triggers swarm initiation. (**A**) Reductionist system for studying neutrophil swarming. Primary human neutrophils are isolated from blood and added to substrates containing patterned ‘targets’ of heat-killed *Candida Albicans* yeast. Neutrophils spread along the yeast targets and initiate propagating self-generated waves of LTB_4_ chemoattractant that recruit additional neutrophils to the targets (neutrophil swarming). Cells are labeled with CalBryte 520AM to assay calcium influx. (**B**) Swarm initiation phases: cells exhibit calcium flux upon engagement of yeast target or when receiving soluble LTB_4_ cue from adjacent neutrophils during swarming waves. These propagating waves of LTB_4_ recruit neutrophils to the target (20x mag, Scale bar: 50 µm). (**C**) Swarm motility 15 minutes after first cell spreading event in **2B**. Target labeled as a grey dot (Scale bar: 100 µm; Radial velocity color scale from −15 to 15 µm/min.). (**D**) High magnification imaging of cells encountering and stretching on the yeast at the patterned target. Neutrophil at origin of swarming wave highlighted with pink asterisk (60x mag, Scale bar: 25 µm). This initiating cell stretches along the single hypha at this yeast target. (**E**) Cell long axis measurements taken along targets patterned with spherical yeast (rounded, smaller) and hyphal (long, filamentous) heat-killed *Candida Albicans* (Spherical yeast engaged neutrophil n = 24; Volunteer N = 3; hyphae engaged neutrophil n = 27; Volunteer N = 4). (**F**) Time measured from first cell contact with yeast to swarm initiation, as measured by first major calcium event (n = 25, Volunteer N = 8).

To better quantify cell morphology preceding initiation events, we repeated these experiments at 60x magnification using both the spherical yeast and the elongated hyphal targets. As previously reported [22, 23], hyphae are potent swarm initiators and induce significant neutrophil stretching prior to initiation. On a representative mixed target containing both yeast clusters and hyphae, the first swarm-initiating wave emerges from the neutrophil stretched most strongly along the hyphae structure **(Figure 2C, Movie 1)**.

Importantly, cell stretch is a consistent feature for all swarm-initiating neutrophils. When encountering either spherical yeast clusters or individual hyphae, neutrophils elongate to almost double their normal length **(Figure 2D)**. While our measurements of pre-initiation cell stretching on hyphae are consistent with previous work [23], it is interesting that similar stretching is also observed for neutrophils that initiate swarms on clusters of spherical yeast. This long axis measurement on clusters is also likely an underestimation, as cells spread in multiple directions across clusters, as opposed to linearly along a single hypha **(Figure 2 – Supplement 1E)**. When stretching along yeast clusters, cells take roughly 8 minutes to initiate swarming **(Figure 2E)**. This time course is only slightly slower than the 4-minute average seen for neutrophils contacting long hyphae targets in a similar *ex vivo* assay [23]. Our data suggest that neutrophil stretching on large targets defines the site of swarm initiation.

### Neutrophil nuclei are strongly deformed following cell contact with yeast clusters

Neutrophils contain segmented, deformable nuclei that serve important roles in their physiology, such as efficient squeezing through small apertures [46]. Because cPLA_2_ is thought to be activated by nuclear stretch, as opposed to overall cell stretch, we next investigated how nuclear morphology evolves during swarm initiation and termination. For precise quantitation of nuclear deformation during swarming, we used Single Objective Light Sheet (SOLS) imaging to gently capture 3D fluorescent volumes during initiation **(Figure 3A)** [47, 48]. This microscope rapidly captures full volumes in a large field of view with minimal phototoxicity, an essential feature for the highly photosensitive process of swarm initiation. We employed a bright, far-red DNA stain to track the morphology of the neutrophil nucleus and used a calcium dye to assay cell engagement with the target and subsequent swarm wave initiation. A representative set of projection stills is shown in **Figure 3B** for a single initiation event, with the initiating cell outlined in the inset (also see **Movie 2**).

**Figure 3.**
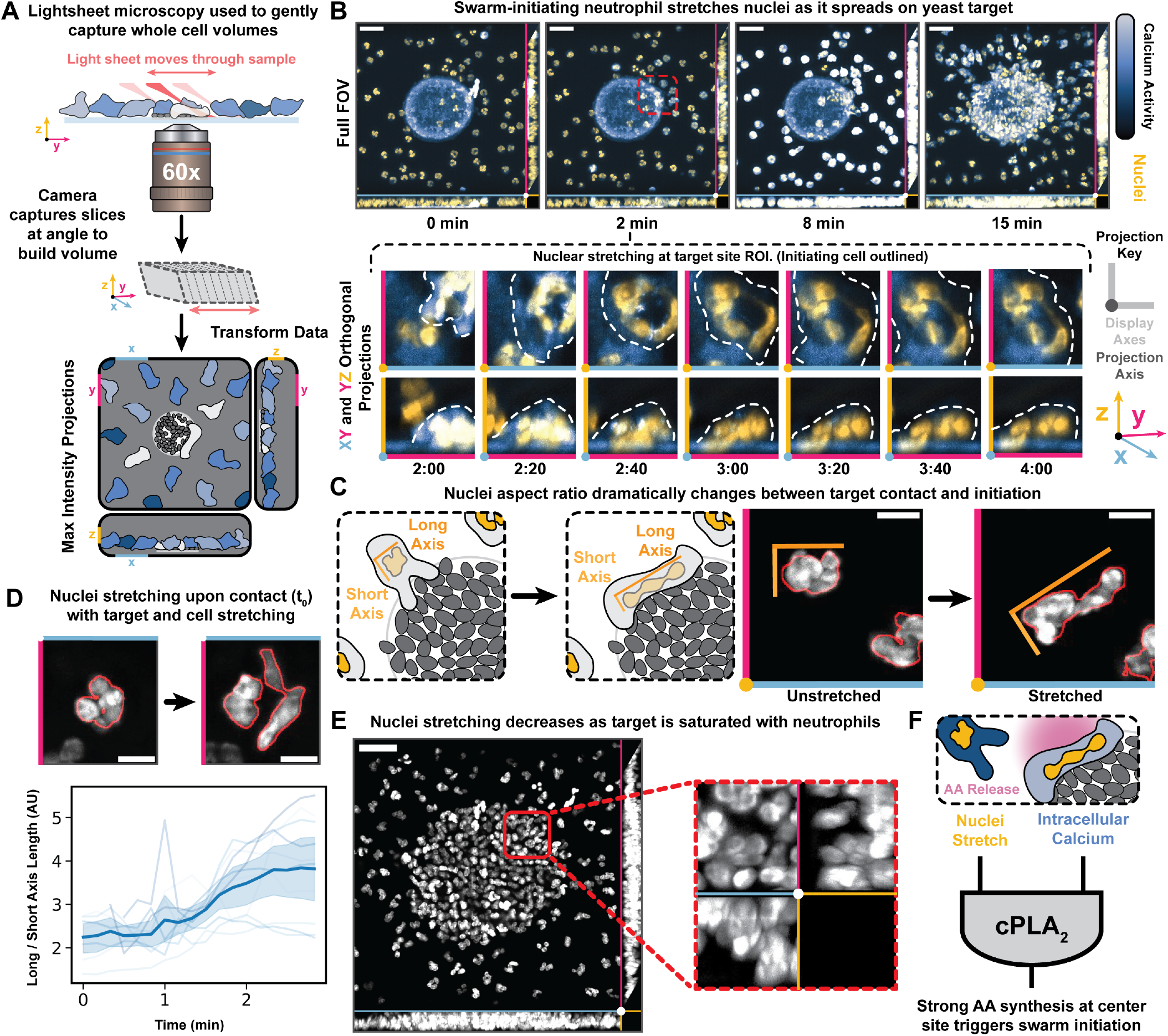
Neutrophil nuclei are strongly deformed following cell contact with yeast clusters. (**A**) Single Objective Light Sheet (SOLS) microscopy was used to capture calcium fluxes (corresponding with propagating LTB_4_ swarming waves), cell movement towards the target, and neutrophil nuclei spreading at the target in 3D with high spatiotemporal resolution. Cells are labeled with CalBryte 520AM and SPY650-DNA. (**B**) Deconvoluted volume rendered as a Max Intensity Projection (MIP) in XY (blue and magneta axis, respectively), XZ (Z axis in yellow), and YZ. Cells engage the target, begin to engulf multiple yeast, and rapidly unfurl their multi-lobed nuclei as they stretch across the target. Inset shows zoomed ROI of “swarm-initiating cell” rendered in XY and YZ. (Scale bar: 25 µm; Insets are 15.5 µm in X and Y dimensions, 13.3 µm in Z.) (**C**) Nuclei stretching can be measured by taking a ratio of the neutrophil nucleus’ long and short axis of the 3D segmented image. Inset shows a cell involved in swarm initiation spreading on the target, resulting in a significant increase in its long-to-short axis nuclear aspect ratio (Scale bars: 5 µm). (**D**) The nuclear aspect ratio (long-over-short axis) measurement of multiple nuclei after neutrophil contacts a cluster of yeast (t = 0) (Scale bars: 5 µm; Nuclei n = 10; Volunteer N = 3). (**E**) At the later phases of swarming, neutrophils crowd on the target, and their nuclei revert to a more compact morphology as cells compete for access to the target. ROI shown is the same as in B where the swarm-initiating cell stretches and triggers a swarm (Scale bar: 25 µm). (**F**) The cPLA_2_ enzyme is an important regulator of neutrophil swarming and generates the AA precursor needed for LTB_4_ production. The coincidence of calcium influx and nuclear stretch activates cPLA_2_. Swarm-initiating neutrophils exhibit both the calcium influx and nuclear stretch needed for cPLA_2_ activation.

For each early cell contact with the target, the nucleus undergoes significant deformation as the cell spreads. By segmenting nuclei and taking the long/short axis measurements in 3D for swarm-initiating cells **(Figure 3C, Supplement 1A)**, we measured how neutrophils unfurl/flatten their uniquely lobed nuclei as they stretch across yeast clusters. Neutrophil nuclei typically begin with a long/short axis ratio of around 2, which increases rapidly following cell stretching and doubles within ∼1 minute **(Figure 3D)**. Some stretching events are so intense that individual lobes of the nucleus are pulled completely apart, and it can be difficult to visualize the small DNA linkages that connect the lobes **(Movie 3, Figure 3 – Supplement 1B)**. As more cells accumulate at the target site, nuclear stretching decreases, and the nuclei return to their more compact morphologies **(Figure 3E, Movie 2)**. If sustained stretch is required for continued cPLA_2_ activation, these observations suggest a potential mechanism for swarm termination.

Together, nuclear stretch and high intracellular calcium in swarm-initiating cells establish the conditions needed to activate cPLA_2_ **(Figure 3F)** [35, 38]. cPLA_2_ is the rate-limiting enzyme that releases arachidonic acid (AA) from cell membranes, and this lipid’s availability is essential for LTB_4_ synthesis and positive feedback [49]. To confirm the importance of this in the swarming context, we next investigated whether the cPLA_2_ enzyme and its downstream product AA play an instructive role in swarm initiation and scaling.

### Neutrophil swarm initiation requires cPLA_2_ activity

To test the functional role of cPLA_2_ activity for swarm initiation, we used the cPLA_2_ inhibitor Pyrrophenone [39, 50] **(Figure 4A)**. Even low levels of this drug (0.5 µM) completely abolish swarm wave initiation from the target **(Figure 4B, Movie 4)**. Notably, patrolling cells continue to migrate normally and rapidly pulse calcium as they spread across a target site (likely due to pathogen recognition receptor engagement [51, 52]) **(Figure 4 – Supplement 1)**. Following spreading, cPLA_2_-inhibited cells fail to initiate propagating calcium waves and large-scale migration toward the target **(Figure 4C, D)**. This defect in collective migration was seen in every repeat of the cPLA_2_ inhibition experiments **(Figure 4E)**.

**Figure 4.**
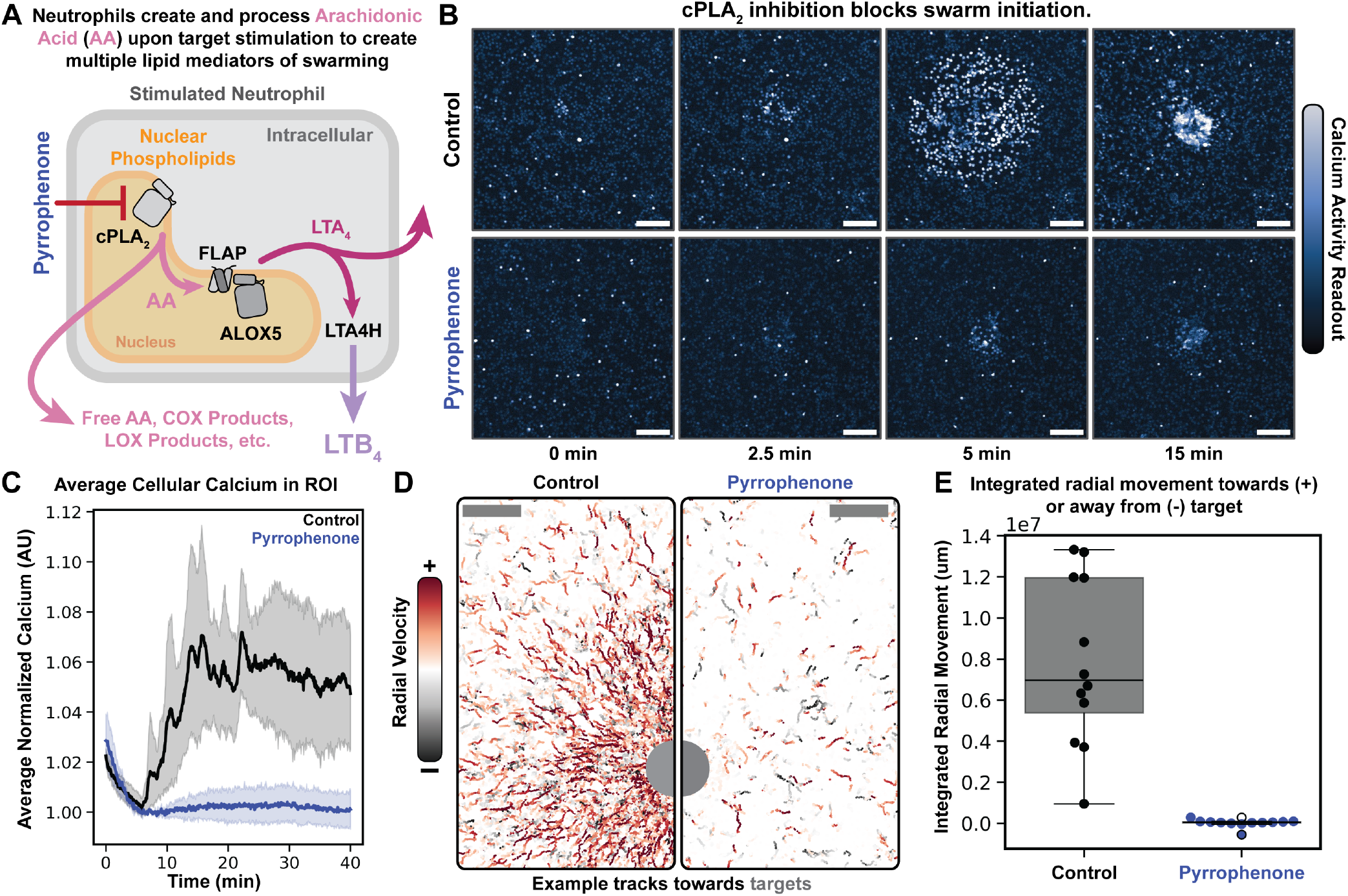
Neutrophil swarm initiation requires cPLA_2_ activity. (**A**) cPLA_2_ frees Arachidonic Acid (AA) from phospholipid stores, and released AA can be converted to LTB_4_, a key swarming mediator, in human neutrophils. (**B**) Pharmacological blockade of cPLA_2_ via Pyrrophenone (0.5 µ_M_) leads to complete inhibition of LTB_4_ swarming wave initiation (Scale bar: 100 µm). (**C**) Average calcium activity across cells over time across target ROIs with and without Pyrrophenone added (Control ROIs = 12; Drug ROIs = 12; Volunteer N = 3). (**D**) Example of neutrophil chemotactic traces towards a target site (labeled with grey dot). Traces show integrated movement 10 minutes after the first cell touches the target, shown in **B** (Scale bar: 100 µm; Radial velocity color scale from −15 to 15 µm/min). (**E**) Integrated radial movement for control and Pyrrophenone-treated samples (sum of radial components of cell tracks for each ROI: positive movement is towards target, negative movement is away from target)(Control ROIs = 12; Drug ROIs = 12; Volunteer N = 3).

Unlike other swarming inhibitors, cPLA_2_ inhibition eliminates calcium activity beyond target engagement, resulting in an inhibition of both local flickering (consistent with blocking diffusive cues from the core) and an inhibition of propagating swarm waves (consistent with blocking LTB_4_ positive feedback relay). In contrast, when neutrophils are treated with a LTB_4_ receptor inhibitor [53], the slow flickering phase of swarm initiation persists, though LTB_4_mediated rapid propagation is inhibited **(Figure 4 – Supplement 2)**. These observations suggest that the slow flickering pre-swarming phase requires the release of AA and could be a readout of subsequent processing into other weakly activating downstream metabolites such as 5-OXO-ETE [54, 55].

### Neutrophils convert free arachidonic acid into LTB_4_ gradients

We hypothesized that calcium influx and stretching of neutrophils on yeast clusters generate a local zone of free AA that fuels the LTB_4_ relay, driving swarm initiation and scaling. If AA is shared locally near the target, exogenous AA gradients should bypass normal swarm triggers. To test this, we plated quiescent neutrophils and exposed them to localized micropipette-delivered AA gradients in the absence of a fungal target. Fluorescent dextran was included to monitor the gradients of AA delivery, and cell responses were quantified using calcium imaging and nuclear-based tracking.

Exogenous AA gradients created via micropipette induce calcium signaling and motility consistent with neutrophil swarm initiation **(Figure 5B, Figure 5 – Supplement 1, Movie 5)**. These responses are not a product of the media flow, nor do they depend on the residual ethanol present in the micropipette **(Figure 5 – Supplement 2, Movie 5)**. For a typical experiment, the fluorescent dextran signal builds rapidly upon applying a positive pressure to the pipette, then quickly (∼10 seconds) elicits a propagating zone of calcium flux in the surrounding cells. Following the calcium flux, cells polarize and migrate towards the pipette, clustering around the centroid of the applied AA gradient. AA is not sensed directly by neutrophils and must be converted to LTB_4_; accordingly, an LTB_4_ receptor inhibitor [53] blocks both the calcium influx and motility responses to exogenous AA gradients **(Figure 5C, D, F, G; Movie 6)**. These waves of activity are similarly blocked by a competitive ALOX5 inhibitor [56, 57] **(Figure 5C, E, F, G; Movie 6)**, again confirming AA conversion to LTB_4_ is a requirement for neutrophil response.

**Figure 5.**
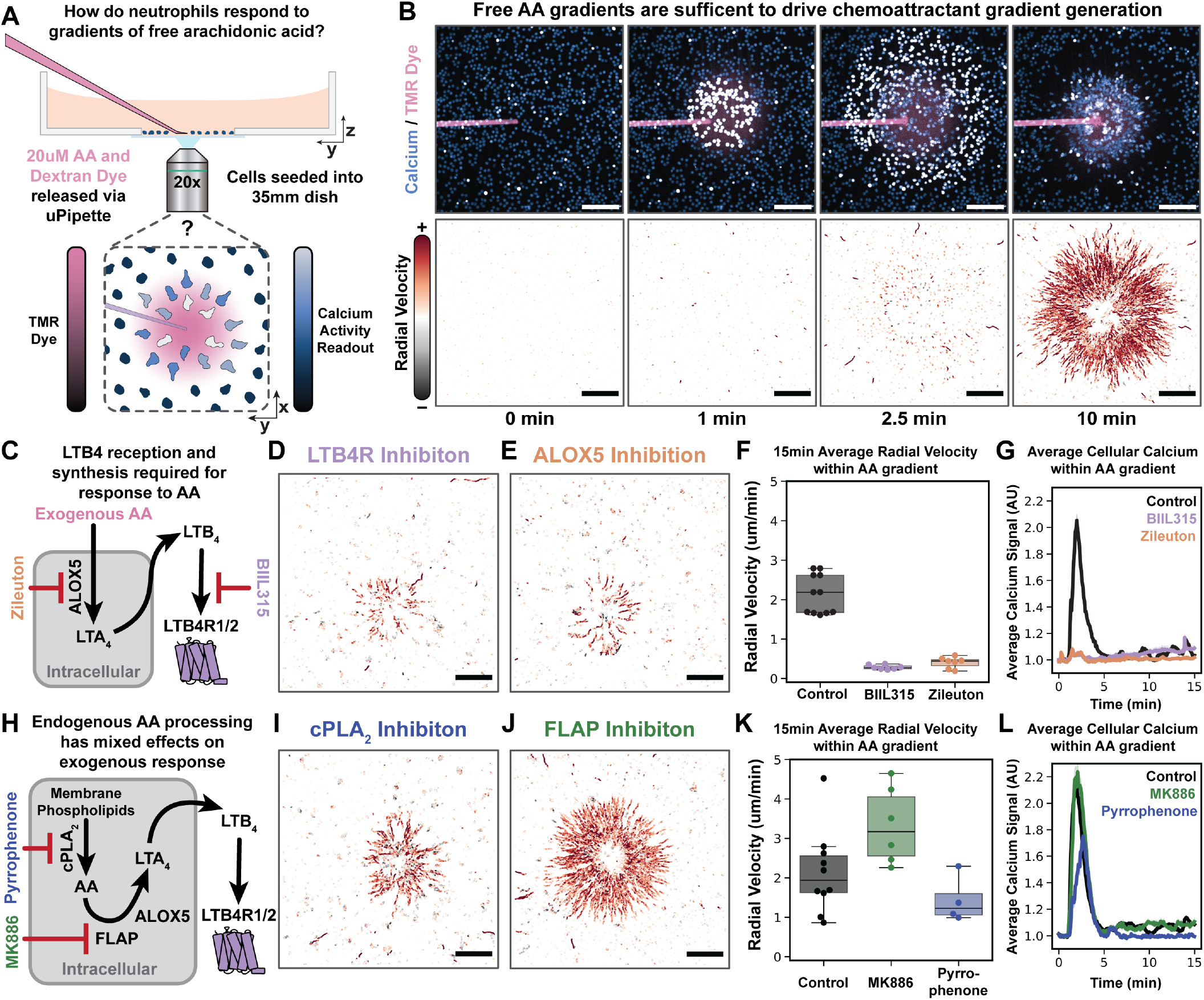
Neutrophils convert free AA into LTB_4_ gradients. (**A**) A gradient of free exogenous AA was delivered to neutrophils via a micropipette loaded with RPMI media containing 20 µ_M_ AA and a 3k dextran conjugated to a tetramethyl rhodamine dye (TMR) (included to assay the shape of the applied gradient). CalBryte 520AM was used to assay calcium responses (indicative of LTB_4_ reception), and Hoechst nuclei tracking was used to assay cell motility. (**B**) Neutrophils rapidly respond to gradients of free AA released from the micropipette with a calcium pulse followed by migration towards the pipette (Scale bar: 100 µm). (**C**) Enzymatic route for conversion of exogenous AA to LTB_4_. Zileuton is used at 50 µ_M_ to block ALOX5-dependent conversion of AA into LTA4, and BIIL315 is used at 1 µ_M_ to block LTB_4_ binding to the LTB_4_ receptor. (**D**) LTB4R inhibition (via BIIL315) strongly impairs neutrophil response to gradients of free AA (Scale bar: 100 µm), indicating that the applied AA is not itself chemotactic. (**E**) ALOX5 enzymatic inhibition (via Zileuton) blocks neutrophils’ ability to respond to gradients of free AA (Scale bar: 100 µm). In conjunction with the LTB4R dependence of the migration, these data indicate that AA must be converted to LTB_4_ by ALOX5 to drive neutrophil chemotaxis. (**F**) Average cell track radial velocity towards the centroid of the dextran gradient taken during a 15min experiment for all cells in frame (Control n = 6; BIIL315 n = 7; Zileuton n = 7; Volunteer N = 4). (**G**) Average cellular calcium signal within applied exogenous AA domain (assayed via dextran gradient) in control, Ziluteon, and BIIL315 conditions (Control n = 6; BIIL315 n = 7; Zileuton n = 7; Volunteer N = 4). (**H**) Enzymatic conversion route of endogenous AA to LTB_4_. Pyrrophenone blocks cPLA_2_ activity (0.5 µ_M_), and MK886 (5 µ_M_) blocks FLAP-based delivery of AA to ALOX5. (**I**) cPLA_2_ inhibition (via Pyrrophenone) mildly impairs neutrophil ability to respond to gradients of free AA (Scale bar: 100 µm); compare to control and also LTB4R or ALOX5 inhibition in **F**. (**J**) FLAP inhibition (via MK886) does not impair neutrophil responses to gradients of free AA (Scale bar: 100 µm). (**K**) Average radial velocity towards the centroid of delivered AA taken during a 15 min experiment for all cells within dextran gradient (Control n = 10; MK886 n = 6; Pyrrophenone n = 4; Volunteer N = 6). (**L**) Average cellular calcium signal within dextran gradient in control, MK886, and Pyrrophenone conditions (Control n = 10; MK886 n = 6; Pyrrophenone n = 4; Volunteer N = 6).

Not all inhibitors of yeast-induced swarming block exogenous AA-mediated migration. Consistent with AA acting downstream of cPLA_2_, the cPLA_2_ inhibitor does not fully suppress AA-induced calcium signaling and motility **(Figure 5H, I, K, L; Movie 7)**. However, its partial effect suggests positive feedback at the level of AA, with cells potentially amplifying exogenous AA stimulation with endogenous AA release, as previously proposed in macrophages [58].

Unexpectedly, inhibition of the adaptor protein FLAP, which delivers AA to ALOX5 for LTB_4_ generation, had little effect. Even at high concentrations of the potent inhibitor MK886, exogenous AA-induced swarming was not reduced and was sometimes slightly enhanced **(Figure 5J, K, L; Movie 7)**. As a positive control for the drug, we showed that a fivefold lower dose of MK886 completely blocks yeast-induced swarming **(Figure 5 – Supplement 2)**. These data suggest that FLAP is dispensable for converting strong, exogenous AA stimuli into LTB_4_.

### Global Arachidonic Acid availability regulates swarm size

In the previous section, we showed that neutrophils convert exogenous AA gradients into local LTB_4_ gradients to enable chemotaxis independent of infection or injury cues. Because target sites may provide multiple sources of AA as many cells stretch across them, we asked whether global AA availability regulates swarm scaling. If AA is a shared metabolite that enables neutrophils to assess the magnitude of infections, then titrating and fixing AA availability should modulate swarm size. To test this, we investigated whether exogenous AA rescues and scales swarming in cPLA_2_-inhibited cells exposed to fungal targets **(Figure 6A)**. To control AA availability, we pre-incubated AA with the lipid carrier human serum albumin (HSA) to stabilize this poorly soluble, oxidation-prone lipid and added it to cells immediately before plating in the target assay.

**Figure 6.**
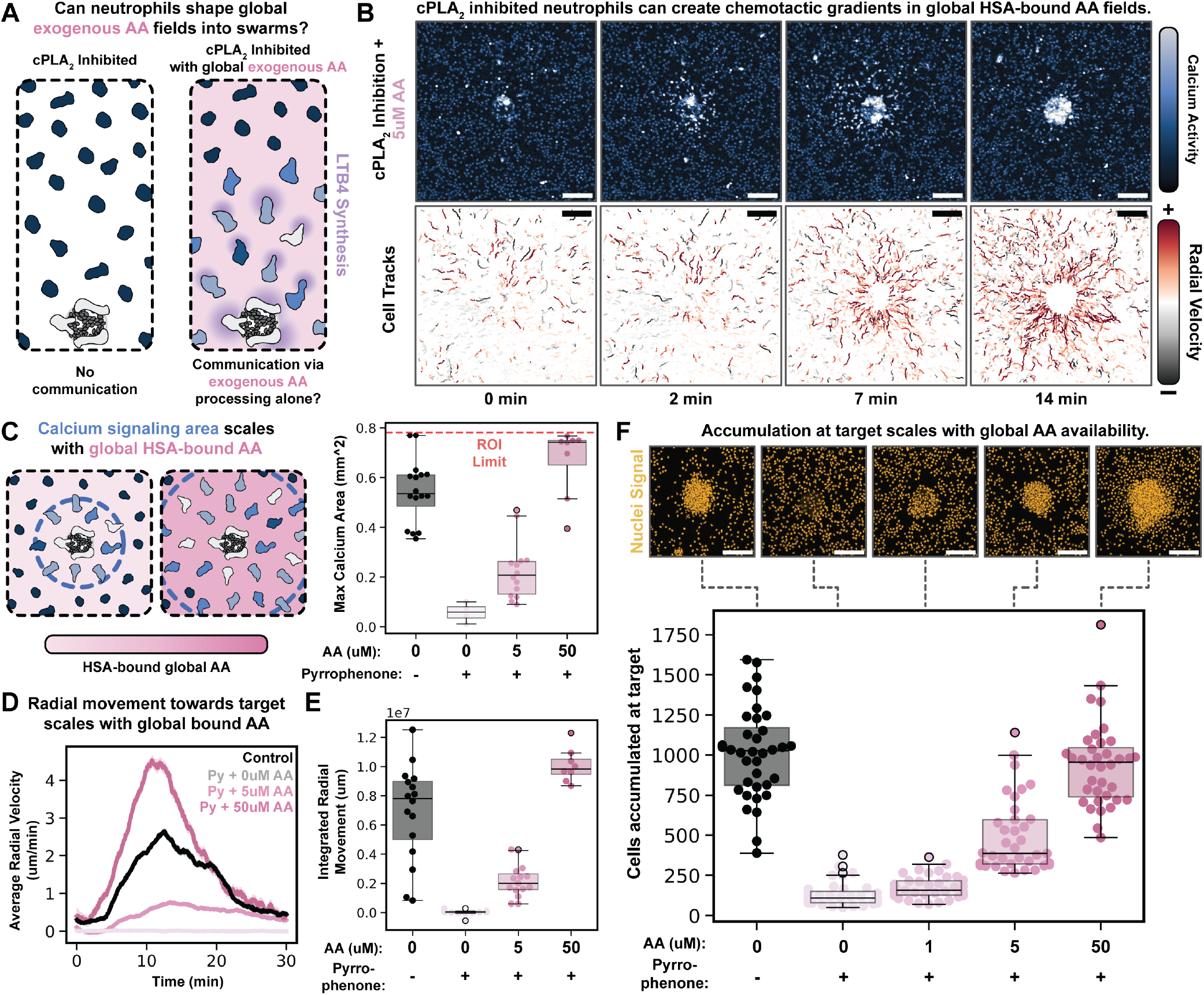
Global arachidonic acid availability regulates swarm size. (**A**) Endogenous cPLA_2_ activity is normally required for neutrophil swarming to yeast targets **(Figure 4)**. Here we tested whether swarming is possible in cPLA_2_ -inhibited cells exposed to uniform exogenous HSA-bound AA. (**B**) Calcium and migration activity of neutrophils responding to a single target ROI in the presence of Pyrrophenone (0.5 µ_M_) and 5 µ_M_ exogenous AA. Swarming is partially recovered under these conditions, and weak calcium activity is observed for cells in the vicinity of the target and is accompanied by motility towards the target (Scale bar: 100 µm). (**C**) Neutrophil swarming waves can be rescued with exogenous AA as the only AA source. Swarming radius scales with exogenous AA concentration, and the highest AA concentrations yield swarming wave radii greater than control cells. (Target ROIs = 42; Volunteer N = 6). (**D**) Radial movement towards the target site over time is averaged across many repeat ROIs and shows that radial migration velocity scales with global AA in the absence of endogenous AA production (Target ROIs = 55; Volunteer N = 6). (**E**) Integrated radial movement towards the target site scales with global AA in the absence of endogenous AA production (Target ROIs = 55; Volunteer N = 6). (**F**) Final accumulation of cells at the arrayed target sites measured 1 hour after adding neutrophils (Target ROIs = 179; Volunteer N = 3). Neutrophil accumulation is strongly inhibited upon cPLA_2_ inhibition, and accumulation at the target can be rescued in a dose-dependent manner by adding increasing amounts of exogenous AA back to the imaging media.

Exogenous AA rescues swarming in neutrophil populations that cannot generate their own, and swarm magnitude scales with global AA availability. Under rescue conditions, neutrophils adhere normally, exhibit calcium flux/stretching at target sites, and rapidly organize into swarms. At 5 µM exogenous AA, cells generate weak calcium waves and limited swarm-based migration **(Figure 6B, Movie 8)**. Increasing exogenous AA 10-fold dramatically increases the radius of swarming waves (assayed via calcium influx), often extending beyond the field of view **(Figure 6C, Figure 6 – Supplement 1A, Movie 8)**. Consistent with unperturbed conditions, these larger swarming waves are accompanied by increased radial migration to-ward the target site **(Figure 6D, E)**.

Endpoint measurements of cell accumulation at target sites show that swarming under cPLA_2_ blockade strongly scales with exogenous AA levels **(Figure 6F)**. Because AA is rate-limiting and must be converted into downstream signals (such as LTB_4_) that directly control swarming, this reflects a layered regulatory system. For targets of a similar size **(Figure 6 – Supplement 1B)** global AA levels primarily determine the number of cells recruited to each site. However, within fixed AA levels, swarm size can still scale with target size **(Figure 6 – Supplement 1C)**, indicating that AA availability is not the only control mechanism for swarm scaling. Together, these data show that AA availability is required for swarm initiation and that AA levels scale the magnitude of swarming responses.

## Discussion

Neutrophil swarming enables a small number of activated cells to rapidly recruit hundreds to thousands of additional cells, a capability that necessitates precise spatial and temporal control. Here, we identify cPLA_2_ activation and AA availability as key regulatory nodes that control both the initiation and scaling of this response. cPLA_2_ activation depends on two inputs: calcium influx and nuclear deformation [35, 42, 59, 60]. Our data suggest that these co-inputs gate swarming in the fungal-triggered context: calcium influx is triggered by yeast contact, whereas nuclear stretch is generated when cells engage large or multiple yeast that cannot be engulfed.

This framework helps explain how receptor-level detection of pathogens alone does not trigger swarming to single spherical yeast cells, but the additional input of nuclear stretch enables robust swarming to yeast clusters or hyphae. Calcium influx and nuclear stretching are uniquely coupled at swarm-initiating sites **(Figures 1,2)**, enabling neutrophils to distinguish between insults that require collective responses versus those that can be resolved individually. The mechanosensitive co-input for sustained AA availability could also provide a mechanism for swarm termination. During later phases of swarming, cells within a swarm maintain high calcium, but their nuclear deformation decreases as other cells crowd the target **(Figure 3)**. Thus, swarm initiation and termination could result from the same mechanosensitive circuit: early nuclear deformation promotes AA production and LTB_4_ amplification, while subsequent loss of deformation attenuates swarm signaling as cells crowd the target site.

Our work also demonstrates a central role for arachidonic acid within the complex swarming regulation circuit. AA availability is necessary **(Figure 4)** and sufficient **(Figure 5)** to initiate neutrophil swarming, and its concentration regulates the magnitude of swarms **(Figure 6)**. Given its role in enabling LTB_4_ positive feedback, one might expect swarming waves to propagate indefinitely in the presence of uniform exogenous AA, but waves still terminate under these conditions **(Figure 6)**. This likely reflects the action of regulatory brakes independent of AA availability, including NADPH oxidase-mediated limitation of LTB_4_ signal relay [8, 61] and GRK-mediated LTB4R desensitization [15]. For a given target size, AA is required for swarming, and swarm magnitude scales with exogenous AA levels **(Figure 6 – Supplement 1B)**. Shared AA among multiple neutrophils engaging a yeast target could enable collective assessment of the magnitude of infection. However, AA alone does not fully determine this response: at fixed AA levels, swarm size still scales with target size **(Figure 6 – Supplement 1C)**. This likely reflects the requirement to convert AA into LTB_4_. This activity is mediated by the strongly calcium-dependent (and potentially stretch-sensitive [37]) enzyme ALOX5, potentially explaining why swarm initiation remains localized to targets even in uniform AA. Together, these findings show that neutrophil swarming is not a binary output but a tunable response, governed by the balance between accelerants such as AA and brakes including NADPH oxidase activity [8, 61] and GRK-mediated receptor desensitization [15].

The mechanosensitive signaling circuit we investigate here may be a general feature of other swarm initiators. For example, calcium influx, nuclear deformation, and cPLA_2_-mediated AA generation are also characteristic features of sterile-injury inflammatory responses [35, 36, 62]. In this context, however, epithelial cells—rather than neutrophils—serve as the primary source of AA, and nuclear deformation is driven by osmotic swelling due to tissue injury instead of pathogen-mediated stretch. Although not yet tested in the sterile injury setting, our data suggest that nuclear stretch-mediated cPLA_2_ activation, AA generation, and shared AA metabolism are unifying features of neutrophil swarming across both injury and infection contexts. A more detailed understanding of how shared lipid metabolic pathways regulate neutrophil swarming will require new experimental approaches. For example, while transcellular sharing of the metabolite intermediate LTA_4_ has been extensively studied in and outside the swarming context [63, 64], genetic tools to study how neutrophils (or neutrophils and other cell types such as platelets or epithelial cells) share AA to coordinate swarming remain underdeveloped and warrant further investigation. Recent tools such as the cPLA_2_ conditional KO mouse [60] and sensors to directly read out LTB_4_ gradients [65] will be indispensable to quantitatively dissect how accelerants and brakes of swarming work together to control when and where swarms occur as well as the magnitude of the resulting response. More broadly, a focus on the AA/LTB_4_ lipid pathway and its regulation will provide key insights needed to better understand, and ultimately control, neutrophil-based inflammation in vivo; with important implications for managing autoinflammatory conditions such as Chronic Granulomatous Disease, rheumatoid arthritis, and Acute Respiratory Distress Syndrome.

## Methods

For a detailed description of all the methods and code used in this work, please see Supplemental Methods.

## Acknowledgments

We thank members of the Weiner lab, particularly Nick Martin for helpful discussion throughout this project. We are deeply indebted to the many blood donors, without whom this project would have been impossible. This work was supported by R35GM118167 (ODW), K22AI163399 (AH), K99GM154115

(HDB), and an AHA predoctoral fellowship (ES). BIIL 315 was kindly provided by Boehringer Ingelheim via its open innovation platform opnMe, available at https://opnme.com. Finally, we would like to thank the Center for Advanced Light Microscopy-Nikon Imaging Center at UCSF for their support in using the custom single-objective light sheet microscope. We are deeply in the debt of Nico Stuurman and Tanner Fadero for helping us operate/debug the SOLS microscope, and thank Kari Herrington and Julia Martin for their tireless efforts maintaining the scope.

## LLM/AI Disclosure

No large language model tools were used in the design of any experiments, nor in any of the writing. No image generation tools were used for any of the graphical design/plotting work. This manuscript was edited with the aid of Grammarly Premium for clarity (advertised to use a blend of GPT4/3.5, Claude, and Llama models). All proposed edits were checked by the authors for accuracy. Large language model tools were used in an assistive capacity for some analysis code written and hand checked by the authors (Microsoft Copilot).

