## Supplemental Figures and Captions for "Arachidonic acid availability controls neutrophil swarm initiation and scaling"

### Supplementary Figures

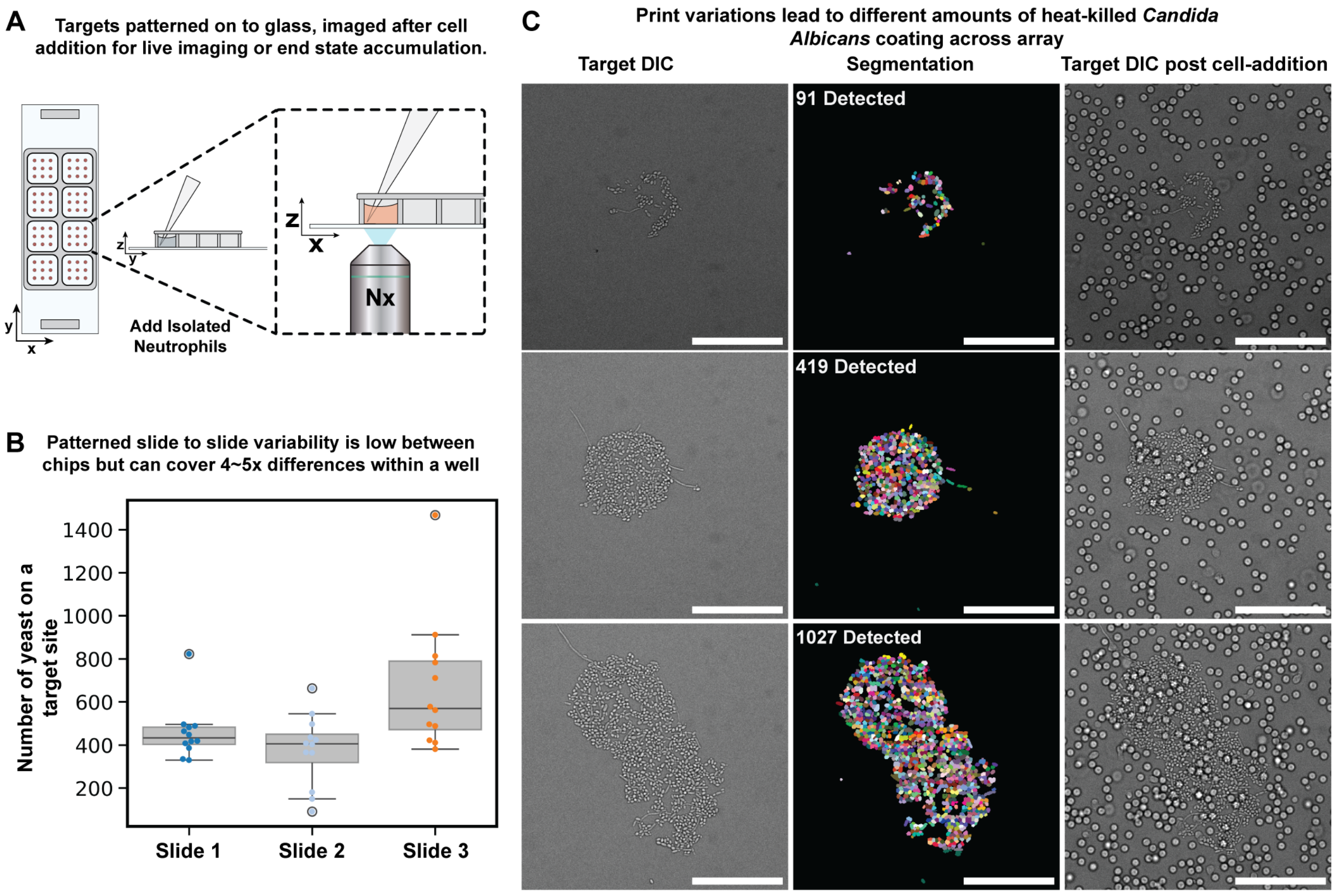


**Figure S1. Figure 1 – Supplement 1**

(**A**) *Ex vivo* assay for neutrophil swarming. Primary human neutrophils are placed on printed targets of heat-killed *Candida albicans*, and neutrophil swarming responses are monitored via timelapse confocal microscopy or by endpoint measurements. (**B**) Slide target patterning is somewhat variable between chips, some receiving more or less yeast, but most variability (4-5x smallest compared to biggest) occurs within a single patterned well (12 targets measured biological replicate). (**C**) Slides are imaged by DIC before the addition of neutrophils, and yeast are segmented and counted using Cellpose to generate labeled masks. Yeast are quantified in a similar manner after neutrophils are added to ensure that the targets remain intact following neutrophil addition. A representative set of example targets with different amounts of printed yeast, imaged both before and after neutrophil addition (Scale bar: 100 µm).

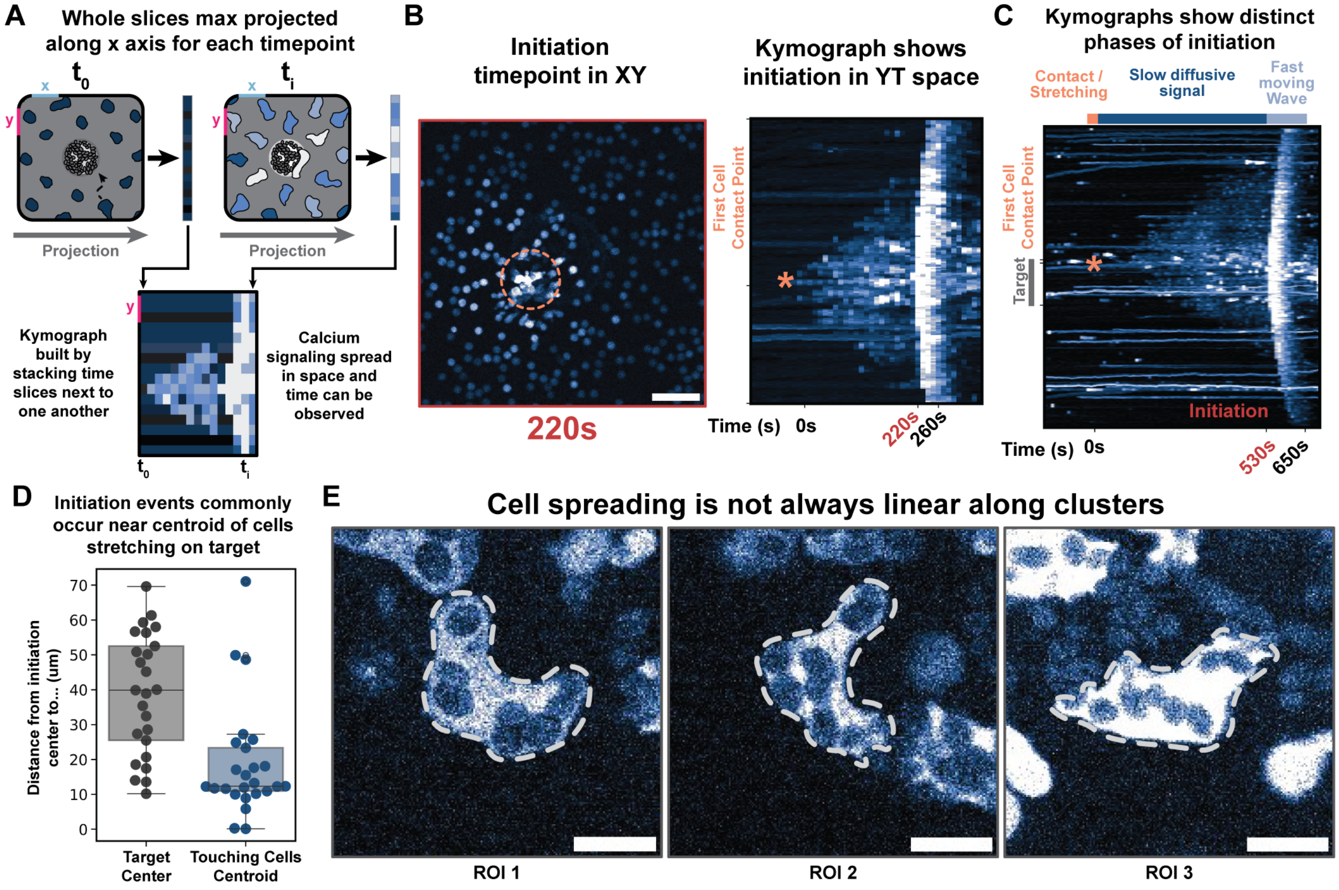


**Figure S2. Figure 2 – Supplement 1**

(**A**) To better visualize the dynamics of swarming waves and their precursors (visualized via calcium influx), kymographs were created by taking the X axis maximum projection of each time slice across the entire ROI. By stacking these linear projections over time, calcium activity can be seen as the vertical spreading of bright areas when read from left to right (forward in time). (**B**) Representative calcium still of **Figure 2B** at the timepoint of initiation. Wave initiating cell is circled, and its place in time and space is marked with an asterisk on the corresponding kymograph for when it first contacted the target site. (**C**) Kymographs reveal three distinct 'phases' of swarm initiation in our system. First a cell (or often a cluster of cells) contacts the target and begins to stretch along it, covering yeast they encounter. Second, a slowly spreading, weakly chemotactic signal can be observed diffusing out from the cell contact sites. After some time, a fast moving, strongly chemotactic wave of activity emerges (usually from the same site), and this begins what classically is known as neutrophil swarming as read out by motility (thus we call this event "swarm initiation" in this work). (**D**) These initiation events often arise closer to the cell or cells in contact with the target site than to the target center itself (Target n = 25; Volunteer N = 8). (**E**) Example ROIs of cells spreading across spherical yeast targets. Cells often contact targets but then generate pseudopods that reach in multiple directions or curve sharply to match yeast geometry, leading to a slight undercount of the long axis measurement (Scale bar: 10 µm).

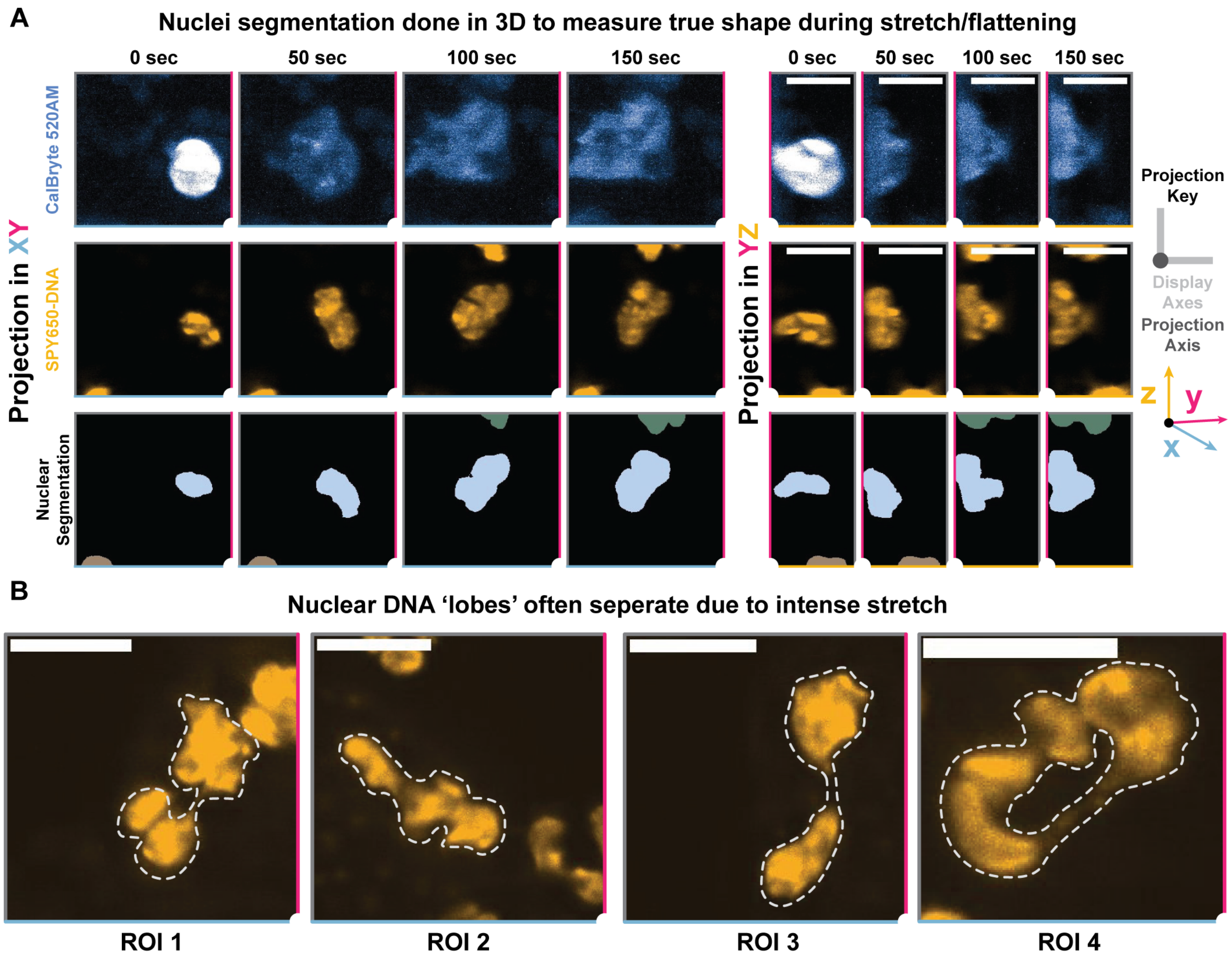


**Figure S3. Figure 3 – Supplement 1**

(**A**) Example XY and YZ projections of cell nuclei for a cell stretching across a fungal target in our light sheet imaging. Cell nuclei are segmented in 3D using an Otsu threshold and labeled as 3D volumes for long/short axis calculations. By segmenting in 3D, we ensure that we are capturing long/short axis changes even during nuclei rotational motion and nucleus flattening events, including the one shown here. Images are cropped from larger ROIs based on where and when a cell first contacts a target, as judged by calcium signaling (Scale Bar: 10 µm). (**B**) Example ROIs of stretching nuclei shown in **Movie 3**. Nuclear lobes often spread to the point of apparent separation or are connected only by thin threads (Scale bar: 10 µm).

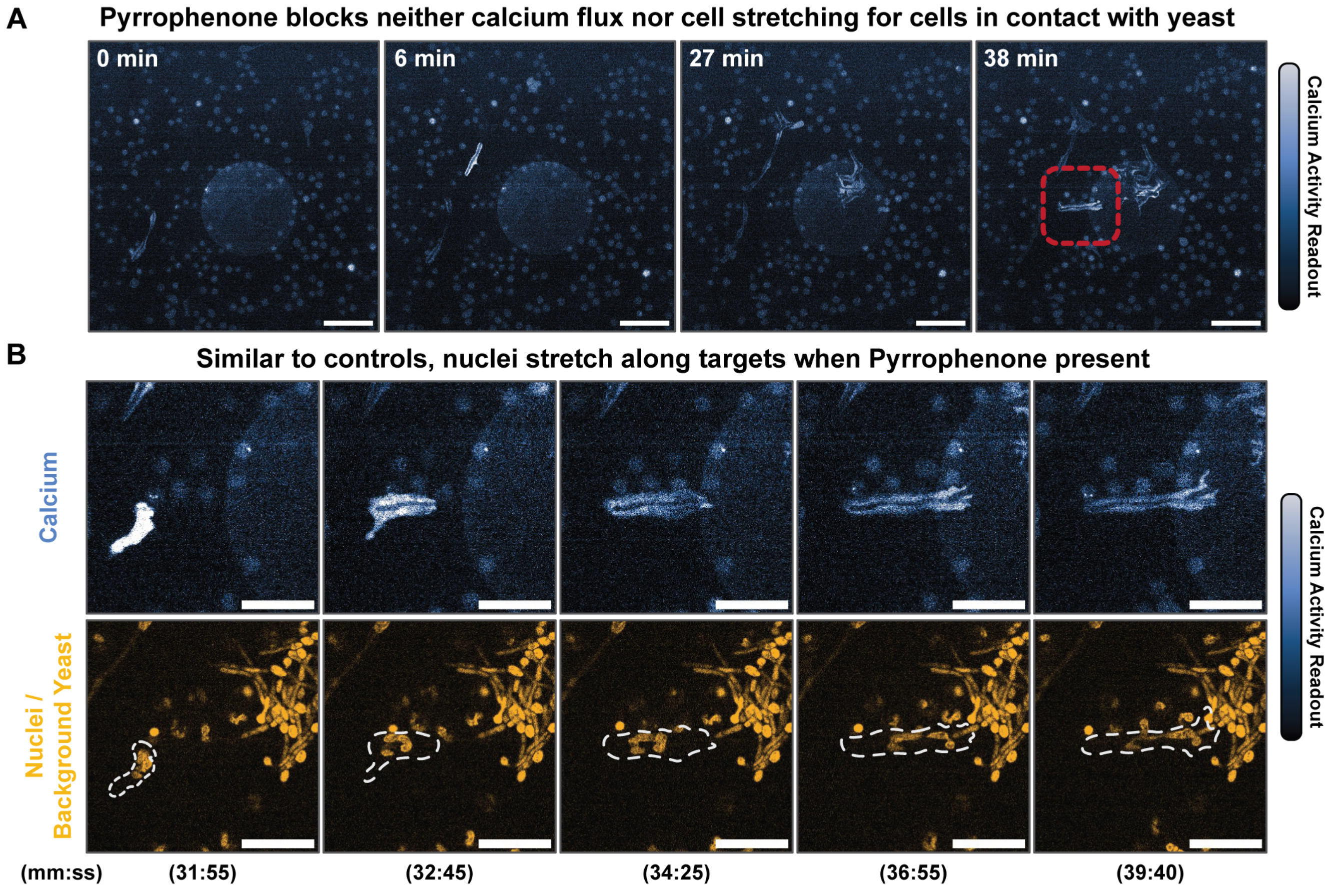


**Figure S4. Figure 4 – Supplement 1**

(**A**) Treatment with 500 nM of Pyrrophenone does not inhibit the ability of neutrophils to settle and spread, patrol randomly until encountering yeast targets, flux calcium upon contact, and stretch along the target yeast. An example 60x ROI is shown with little to no propagation of signals from the cell on the target to cells off the target, as assayed by calcium. There is also a lack of directed movement of cells towards the target after 40 minutes (Scale bar: 50 µm). (**B**) Zoomed inset ROI from **A**. Neutrophil encounters hyphae and elongates along it, stretching the nucleus (labeled with SPY-650DNA) of the cell without initiating a swarm following Pyrrophenone treatment. Some hyphal yeast are auto fluorescent in far-red and show up in the nuclear channel (Scale bar: 25 µm).

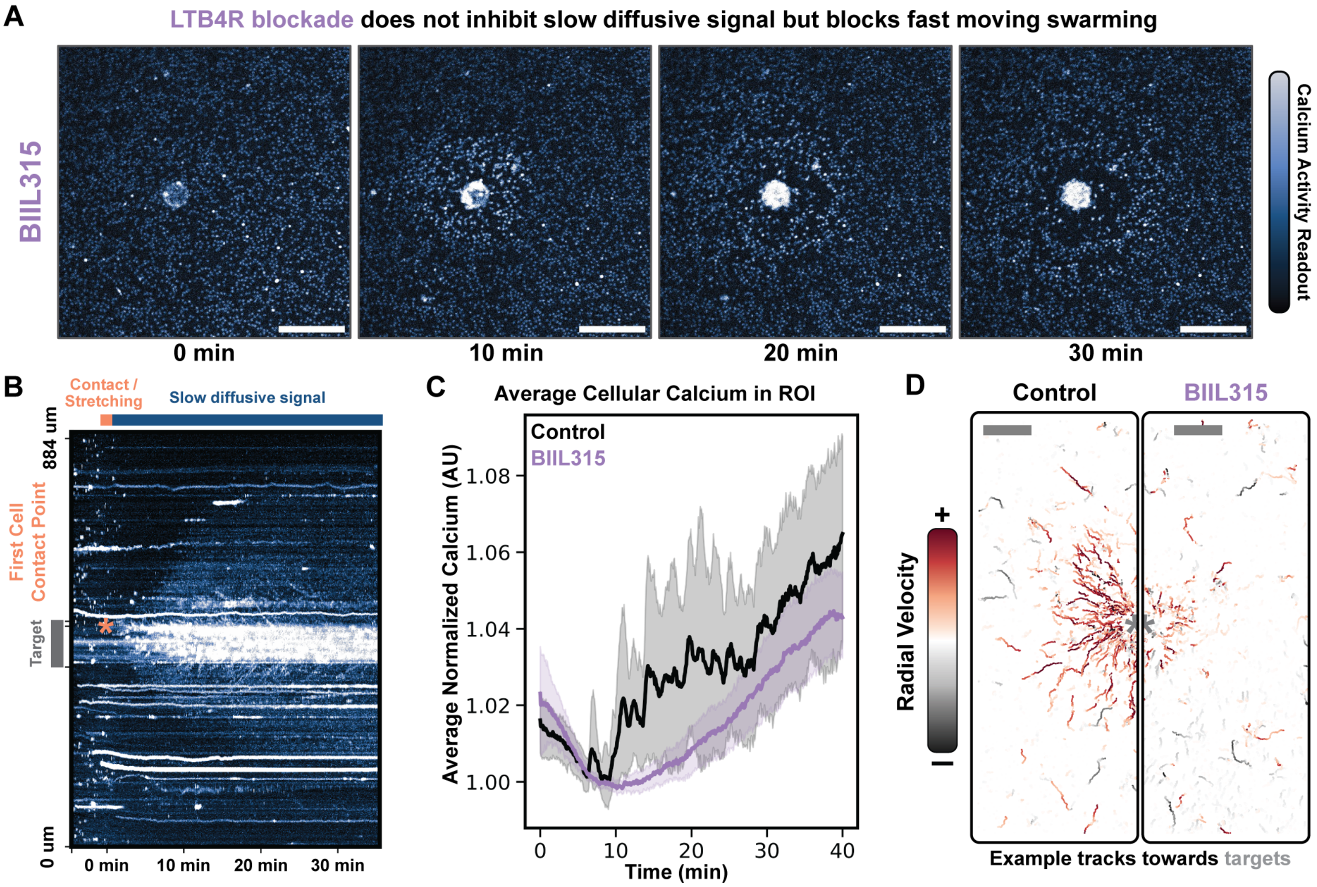


**Figure S5. Figure 4 – Supplement 2**

(**A**) $LTB_4$ receptor blockade via 1 µM BIIL315 blocks fast moving waves and thus swarm initiation but does not inhibit slow-moving diffusive signals. Example time course is shown of cells responding to a target while drug is present (Scale bar: 200 µm). (**B**) Kymograph of **A** showing how slow diffusive signals continue to fan out from cells at the center site, even though no large, fast-moving waves emerge. (**C**) Average cellular calcium of multiple ROIs averaged across many runs (Control n = 11; Drug n = 10; Volunteer N = 3). (**D**) Example radial velocity plot for a given volunteer control and corresponding drug treated ROI. LTB4R inhibition inhibits fast-propagating swarming waves and restricts chemotactic movement to the cells that are very close to the target (Scale bar: 100 µm).

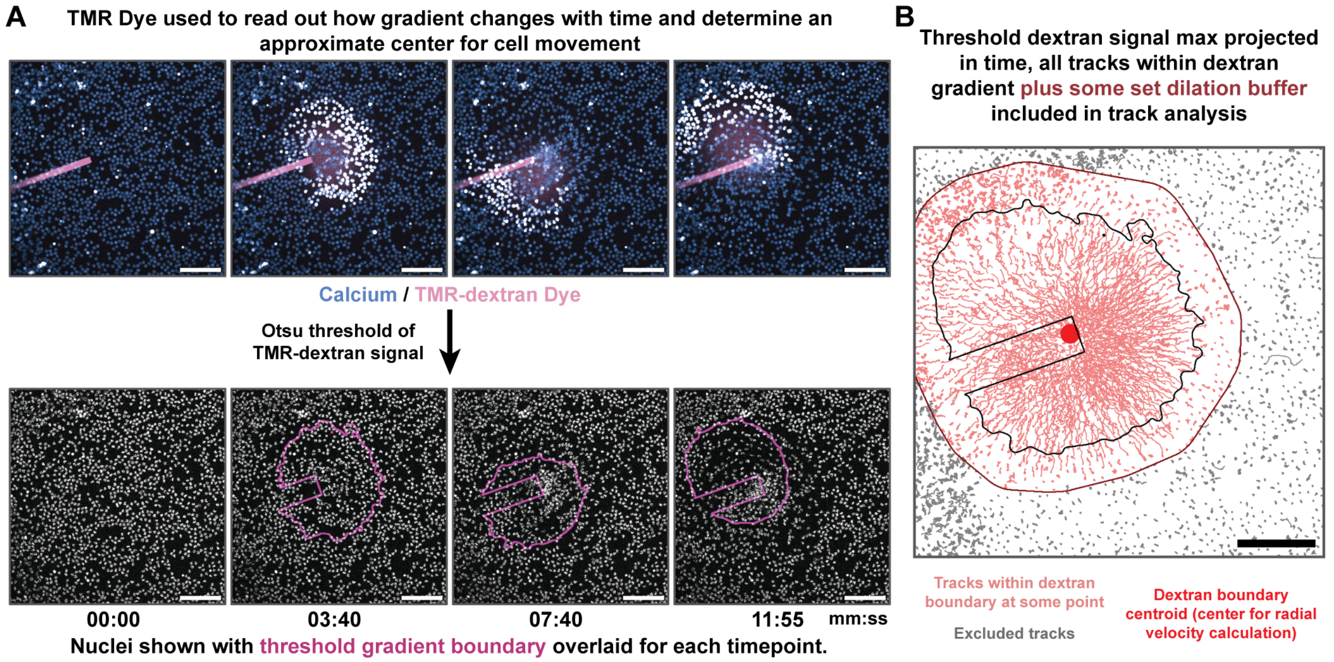


**Figure S6. Figure 5 – Supplement 1**

(**A**) Neutrophil response to pipette read out by analyzing behavior within an estimated AA boundary (assayed via TMR-dextran signal). To establish this boundary, the pipette is first algorithmically detected by its intense brightness, and a box is drawn around the pipette to exclude it from further boundary calculations. The remaining TMR signal is binarized with an Otsu threshold calculated for the whole video and applied to each frame (Scale bar: 100 µm). (**B**) To determine which cell tracks to analyze for calcium and movement, the max projection of the binarized TMR signal is taken across all timepoints to establish the locations subjected to AA. This boundary is dilated to provide a generous buffer for cell tracking, and the centroid of this shape is used as the center towards which radial velocities are calculated (Scale bar: 100 µm).

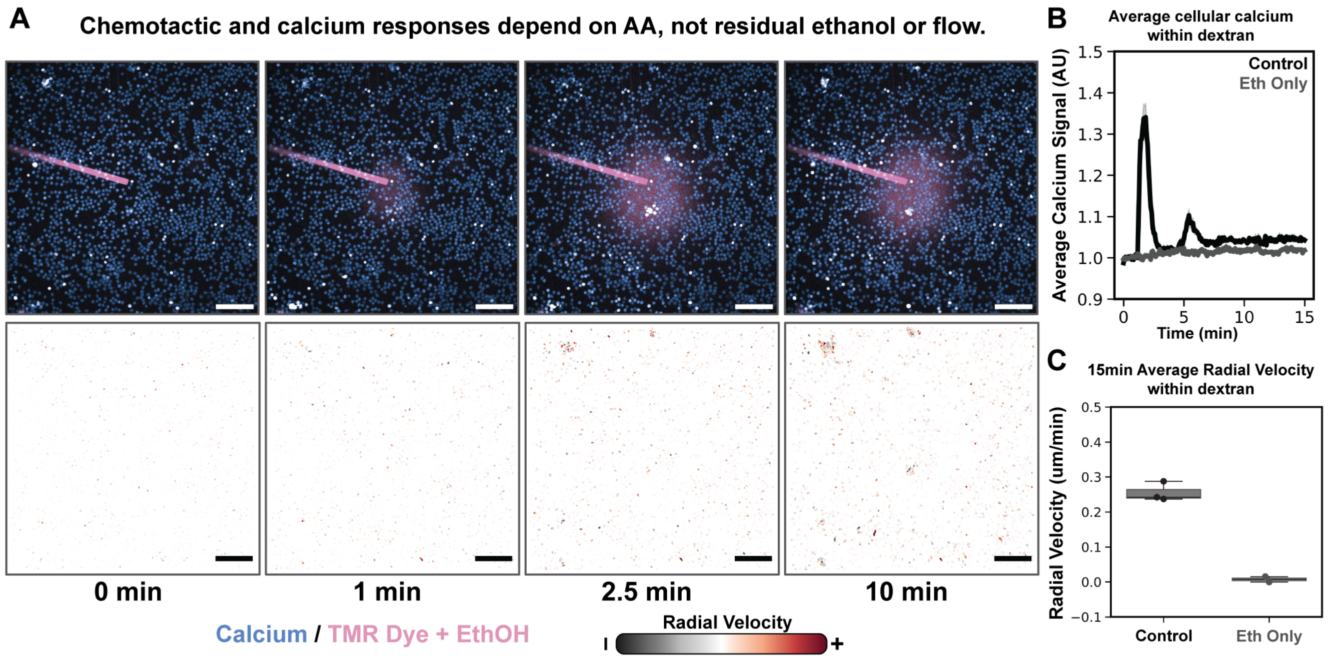


**Figure S7. Figure 5 – Supplement 2**

(**A**) Neutrophil response to pipette depends on neither the flow generated by the pipette nor the residual ethanol present in our free AA mixes. Exogenous, free AA is required for cells to respond and migrate towards the micropipette (Scale bar: 100 µm). (**B**) Average calcium response for a matched 20 µM AA control and ethanol only run (Experimental n = 2 each condition; Volunteer N = 1). (**C**) Resulting motility for matched 20 µM AA control and ethanol only micropipette experiments. Ethanol and TMR dye alone elicit neither calcium signaling nor migration to the pipette (Experimental n = 2 each condition; Volunteer N = 1).

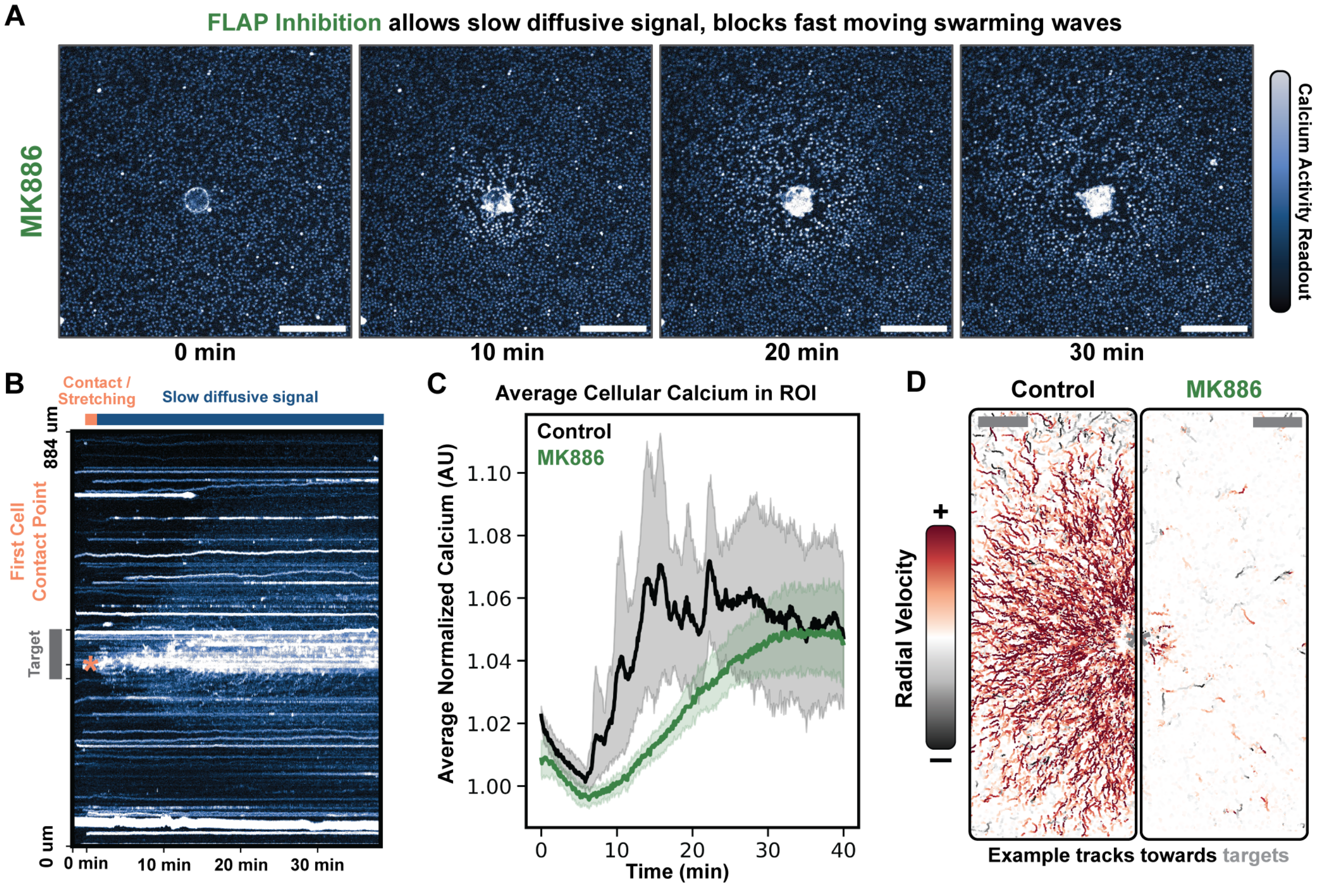


**Figure S8. Figure 5 – Supplement 3**

(**A**) FLAP inhibition blockade via 1 µM MK886 blocks fast moving waves and thus swarm initiation, but not slow-moving diffusive signals. Example time course is shown for cells responding to a target while drug is present (Scale bar: 200 µm). (**B**) Kymograph of **A** showing slow diffusive signal generation from cells at the center site, even though no large, fast-moving waves emerge. (**C**) Cellular calcium of multiple ROIs averaged across many runs (Control n = 12; Drug n = 12; Volunteer N = 3). (**D**) Example radial velocity plot for a given volunteer for control versus corresponding drug treated ROI. While FLAP inhibition limits swarming to yeast targets at 1 µM concentration, 5 µM of drug fails to block neutrophils' ability to convert exogenous AA gradients into chemotactic signals in a micropipette context **(Figure 5J,K,L)**. These data reveal an underappreciated role of exogenous AA in bypassing canonical pathways that regulate $LTB_4$ synthesis (Scale bar: 100 µm).

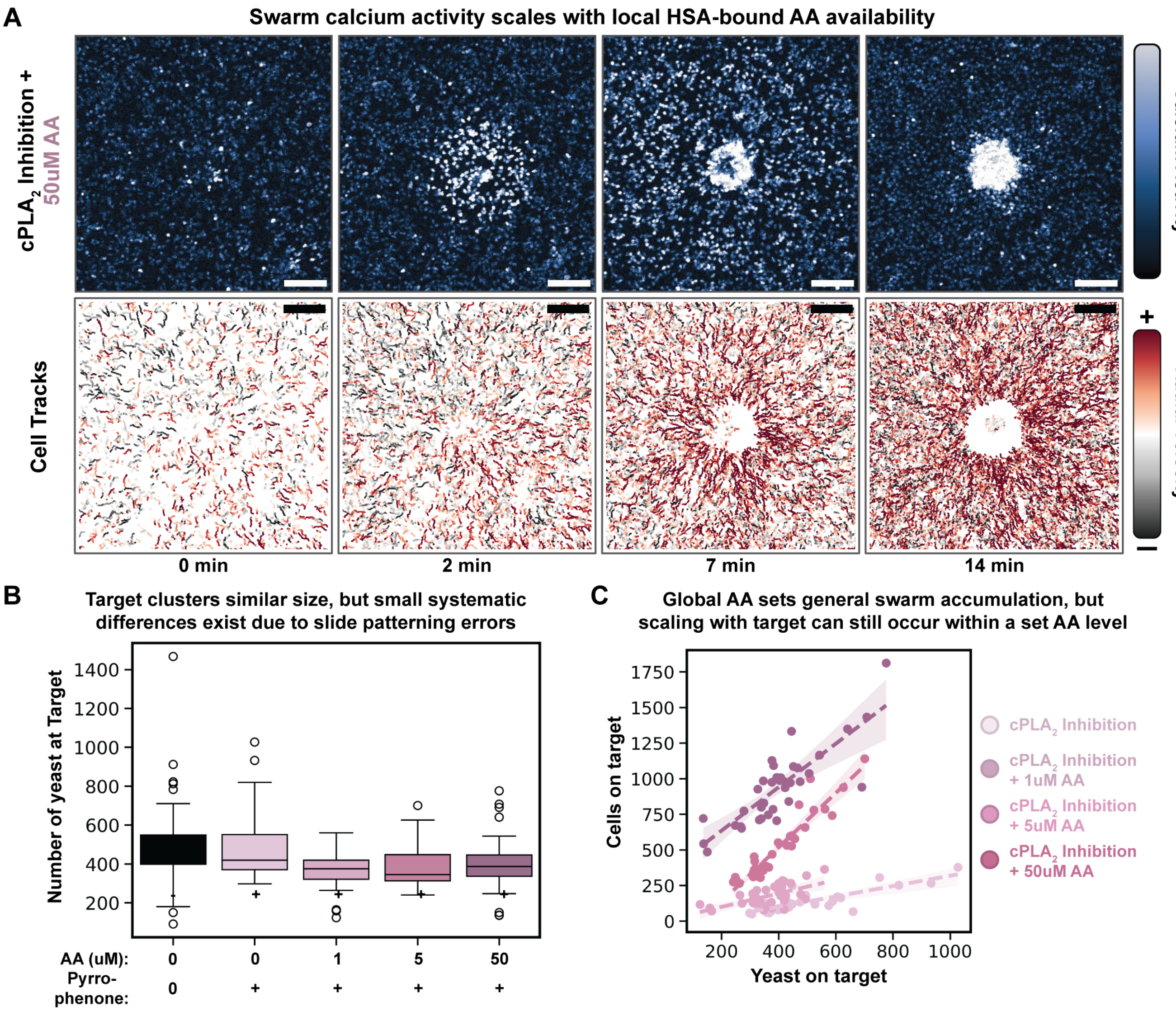


**Figure S9. Figure 6 – Supplement 1**

(**A**) Neutrophil swarms under Pyrrophenone (0.5 μM) endogenous AA synthesis blockade supplemented with 50 μM exogenous AA. Cells on target areas trigger strong calcium waves that sweep out of ROI view and create very strong chemotaxis into the target center (Scale bar: 100 μm). (**B**) Target clusters systematically vary by condition due to unforeseen biases in the coating of the slides during manufacturing. Regardless of these small systematic errors, the resulting swarming trends show a strong dependence on global AA availability (Target n = 179). (**C**) While global AA sets the general swarm size, other regulatory mechanisms are layered into this system to enable some scaling to the target site size at a fixed amount of AA. (Target n = 179; Volunteer N = 3).

### Supplementary Video Captions

**Movie 1: Human Neutrophil swarming *ex vivo*.**
Confocal imaging of neutrophils swarming to spots of yeast clusters and hyphae, as read out by intracellular calcium dyes. Increased brightness (blue to white color changes) indicates calcium influx and cell activation. A series of example movies are shown that make up the stills for **Figure 2B** and **2C**. Neutrophils can be seen stretching across multiple yeast clusters before swarm initiation for each event. Targets that include only yeast clusters and only hyphae are shown for videos that were quantified in **Figure 2E**. All scale bars are given in video title cards.

**Movie 2: Human neutrophil target engagement imaged via Single Objective Light-sheet Microscopy.**
3D light-sheet imaging of neutrophils swarming to spots of yeast clusters and hyphae, as read out by intracellular calcium dyes and simultaneous imaging of nuclear morphology. Increased brightness (blue to white color changes) indicates calcium influx and cell activation. Nuclei labeled via SPY-DNA dye and represented in gold. A series of example movies are shown that make up the stills for **Figure 3B** and the zoomed inset for **Figure 3B**. Neutrophils stretching across yeast clusters can be seen to strongly stretch their nuclei. At later time points in imaging, stretched nuclei contract again as neutrophils crowd the target site. All scale bars are given in video title cards.

**Movie 3: Examples of nuclear stretching upon neutrophil-yeast engagement**
Three example light-sheet ROIs of neutrophil nucleus stretching upon cell contact with yeast clusters. Nuclei are labeled via SPY-DNA dye and represented in greyscale. Nuclear segments can be seen stretching to such an extent that individual DNA lobes separate, and little fluorescent signal can be seen between lobes. All scale bars are given in video title cards.

**Movie 4: Neutrophil swarming is inhibited by the cPLA$_2$ inhibitor, Pyrrophenone**
Confocal imaging of neutrophils interacting with yeast under Pyrrophenone treatment (all videos at 500 nM concentration). Neutrophil swarm activity is completely ablated, though cells can still move and interact with the yeast target. Neutrophils are shown interacting with hyphae during Pyrrophenone treatment. Nuclei are labeled via a SPY-DNA dye, and some yeast also exhibit background autofluorescence in the nuclei fluorescence channel. Neutrophils can be seen stretching normally along hyphae during Pyrrophenone treatment, but no swarming results. All scale bars are given in video title cards.

**Movie 5: Arachidonic acid is sufficient to drive swarm signaling**
Confocal imaging of neutrophils responding to 20 µM of arachidonic acid released locally via micropipette. Increased brightness (blue to white color changes) indicates calcium influx and cell activation. Approximate location of arachidonic acid visualized with a fluorescent dextran tracer included in the pipette. Neutrophils respond to the released arachidonic acid with an approximate 5-10 second delay. Upon activation, neutrophils flux calcium and migrate towards the centroid of the arachidonic acid gradient. An example ROI is shown first that corresponds to **Figure 5B**. Activation depends on arachidonic acid release from the pipette, as the last video shows a lack of neutrophil activation when the pipette releases dextran and carrier ethanol alone. All scale bars are given in video title cards.

**Movie 6: LTB$_4$ signaling is required for neutrophil response to exogenous AA**
Confocal imaging of neutrophils responding to 20 µM of arachidonic acid released locally via micropipette in the presence of various pharmacological inhibitors that are present in both the cell media and the pipette. Increased brightness (blue to white color changes) indicates calcium influx and cell activation. Blocking LTB$_4$ reception via 1 µM of the LTB4R inhibitor BIIL315 dramatically reduces the ability of neutrophils to respond to the released AA. Similarly, 50 µM of the ALOX5 inhibitor Zileuton reduces both calcium influx and motility in response to released AA. All scale bars are given in video title cards.

**Movie 7: The effect of FLAP and cPLA$_2$ inhibition on neutrophil response to exogenous AA**
Confocal imaging of neutrophils responding to 20 µM of arachidonic acid released locally via micropipette in the presence of various pharmacological inhibitors that are present in both the cell media and the pipette. Increased brightness (blue to white color changes) indicates calcium influx and cell activation. Blocking FLAP activity via 5 µM of MK886 does not block neutrophil response to exogenously released AA. Blocking cPLA$_2$ activity via 0.5 µM of Pyrrophenone partially blocks neutrophil response to exogenously released AA. All scale bars are given in video title cards.

**Movie 8: Global AA availability modulates swarm response to yeast clusters**
Confocal imaging of neutrophils interacting with yeast under Pyrrophenone treatment with and without exogenous arachidonic acid (all videos at 0.5 µM Pyrrophenone concentration). Increased brightness (blue to white color changes) indicates calcium influx and cell activation. Swarm activity is rescued by the addition of exogenous arachidonic acid for cells that are unable to synthesize their own AA endogenously. All scale bars are given in video title cards. Presenting cells with 5 µM of AA elicits a mild rescue of swarm behaviors, while 50 µM of AA enables large swarm waves that recruit neutrophils from beyond the microscope field of view. All scale bars are given in video title cards.
