## Supplemental Methods for "Arachidonic acid availability controls neutrophil swarm initiation and scaling"

### Materials and Methods for Submission

**E. Strickland et al. 2026.**

**Reagents Table**

| REAGENTS | SOURCE | IDENTIFIER |
|---|---|---|
| **Chemicals, peptides, and recombinant proteins** | | |
| RPMI (w/o Phenol Red, L-Glutamate) | Gibco | 11835030 |
| 1M HEPES | Gibco | 15630080 |
| Human Serum Albumin | Sigma | A5843 |
| Hoechst 3334 | Invitrogen | H3570 |
| CalBryte 520 AM | AAT Bioquest | 20650 |
| SPY-650 DNA | Cytoskeleton | CY-SC501 |
| Heat inactivated FBS | Gibco | 16140-071 |
| Poly-l-lysine | Sigma-Aldrich | P8920-100ml |
| ZETAG 8185 | Solenic | N/A |
| Porcine Fibronectin | Lab-Generated | Procol based on publication[1] |
| DPBS (no Ca/Mg) | Gibco | 14190-144 |
| BIIL315 | Boehringer Ingelheim | N/A |
| MK886 (10mM * 1mL in DMSO) | MedChemExpress | HY-14166 |
| Pyrrophenone – 1mg Solid | MedChemExpress | HY-111376 |
| Zileuton – Solution 10mM * 1mL in DMSO | MedChemExpress | HY-14164 |
| Arachidonic Acid – 100mg | Cayman Chemical | 90010 |
| Dextran, Tetramethylrhodamine, 3000 MW, Anionic | ThermoFisher | D3307 |
| Anhydrous DMSO | Invitrogen | D12345 |
| **Critical commercial assays** | | |
| Stemcell EasySep Direct Human Neutrophil Isolation Kit | StemCell Technologies | 19666 |
| Stemcell Big Easy Magnet | StemCell Technologies | 18001 |
| Stemcell Easy50 Magnet | StemCell Technologies | 18002 |
| **Software and algorithms** | | |
| Fiji/ImageJ2 (v2.14.0/1.54f) | N/A | https://fiji.sc/ |
| Python/MiniConda | N/A | https://anaconda.org/ |
| ARCOS | N/A | https://arcos.gitbook.io/home/ |
| Stardist | N/A | https://github.com/stardist/stardist |
| **Other** | | |
| .22um Seriflip Filter | Millipore Sigma | SE1M179M6 |
| 23-gauge butterfly needle collection set | BD | 23-021-022 |
| Vacutainer EDTA tubes | BD | 366643 |
| 1.5H Glass Coverslip | Ibidi | 10812 |
| 8-well sticky-Slide | Ibidi | 80828 |
| Mattek 35 mm Dish \| No. 1.5 Coverslip | Mattek | P35G-1.5-14-C |
| Spermatic Injection Pipettes for Piezo ID:9.0 um / 35 / 10 (Custom ordered for polished tips) | Sunlight Medical | SIC-90F-35 |
| Fisherbrand LowRetention 1.5mL microcentrifuge tubes | Fisher | 02681320 |
| Hellmanex III | Hellma Analytics | |
| Vials, screw top with solid green Thermoset cap with PTFE liner | Sigma Aldrich | 27001-U |

*All reagents listed here were used in this protocol and ordered with the catalogue numbers given.

**Study Participant Details:**

Healthy blood specimens from healthy volunteers were obtained with informed consent according to the institutional review board-approved study protocol at the University of California, San Francisco (Study #21-35147). Volunteers were informed not to take ibuprofen or acetaminophen within 48 h and aspirin within 7 days of blood draw. Volunteers were asked to consume no more than one alcoholic drink the night before blood draw. Volunteer demographics such as age and sex are provided in the final section of this methods section. Informed consent was obtained for use of human tissues in research.

**Method Details:**

Neutrophil Isolation Protocol:
Imaging media was first prepared with RPMI (w/o Phenol Red, 25mM HEPES, L-Glutamate) and 0.4% Human Serum Albumin (HSA) (Sigma). HSA was added directly to RPMI and then centrifuged at 500 x g for 5 min until fully dissolved. The mix was then filtered with a 0.22 µm Steriflip filter (Millipore Sigma) before further use. Imaging media was always prepared fresh on the same day of imaging.

Fresh samples of peripheral blood (2 tubes, 7 mL each) from healthy adult volunteers were collected via a BD 23-gauge butterfly needle collection set into 10 mL BD Vacutainer EDTA tubes. Blood was kept on a shaker at minimum setting and utilized within 2 hours of the draw. Neutrophils were isolated using the Stemcell EasySep Direct Human Neutrophil Isolation Kit (Stemcell) with the BigEasy magnet (Stemcell) according to the manufacturer's protocol.

For most experiments, isolated neutrophils were spun down at 200 x g for 5 min and then resuspended in a dye media consisting of imaging media plus 5 µg/ml Hoechst 3334 (Invitrogen), and 1 µM CalBryte 520 AM (AAT Bioquest). **For light sheet experiments**, no Hoechst was added to the dye media (only CalBryte520); please see below for modification to this step. This cell suspension was incubated at room temperature in the dark for 15 min, and then spun down at 200 x g for 5 min. The dye medium was aspirated and replaced with an amount of cell culture media of RPMI (Gibco) with 10% heat-inactivated FBS (Gibco) needed to achieve a final cell density at or below 1 x $10^6$ cells/mL. Purified neutrophils were then kept in polystyrene T25 flasks at 37°C in a 5% CO2 environment until imaging. Cells were allowed to incubate for at least one hour before imaging began and not more than 5 hours after isolation. Allowing the cells to rest in culture before imaging helped ensure that $Ca^{2+}$ signaling was more consistent and less noisy across volunteers.

Neutrophil Swarming Chip Manufacturing Protocol:
Swarming arrays of *C. albicans* were prepared as described previously[2,3]. Briefly, we used a microprinting platform (Picospotter PolyPico Galway, Ireland) to print a solution of 0.1% poly-l-lysine (Sigma-Aldrich) with ZETAG 8185. Coverslips were washed in ethanol and dried prior to printing to ensure a clean surface. For experiments, we printed arrays with 1.0 mm spacing in an 8 well format on full sized No. 1.5H glass coverslips (Ibidi). Coverslips were dried (often overnight) at 37°C and then left at room temperature until the next step could be performed. To attach an infection-like material to these arrays, 8-well sticky-Slide attachments (Ibidi) were overlaid on the printed arrays. **For spherical yeast only cluster targets**, an overnight culture of live C. albicans yeast was heat killed at 90°C for 20 minutes before being washed and re-suspended in dH2O, then 750 µL of this suspension was added to each well and incubated for 5 minutes. **For hyphal targets**, live *C. albicans* yeast from an overnight culture were transferred to RPMI and cultured again overnight to induce hyphal growth. The next day hyphae were concentrated and were then heat killed, washed and added to each well as outlined for yeast. Following incubation, wells were thoroughly washed out with $dH_2O$ to remove unbound targets from the glass surface. Wells were screened to ensure appropriate patterning of targets onto the spots with minimal non-specific binding before use. The well attachment was removed, and the coverslips were stored at 4°C until ready for use.

When preparing to use a patterned coverslip for a swarm assay, the coverslip was first re-inspected for coating and target integrity. An Ibidi 8 well sticky-slide was pressed firmly on to the slide, and a pipette tip was run along the bottom to ensure a proper seal was formed. The coverslip-well combo was then incubated in a 37°C oven overnight. Next a 200 µL mixture of imaging media plus 16 µg/ml fibronectin (lab prepared, porcine origin) was pipetted into each well for use each day. The slide was then incubated for 30 min at 37°C and washed 3x with 200 µL/well of PBS (-/- Ca/Mg). The final wash of PBS was left on the well until imaging.

General Neutrophil Swarming Live Imaging Protocol:

Cells were imaged as follows for all confocal swarming experiments for both spherical yeast and hyphal yeast targets (See below for lightsheet and micropipette setups). Neutrophils in culture were taken and placed into Fisherbrand LowRetention 1.5 mL microcentrifuge tubes and spun down at 200xg for 5 min. Cells were resuspended at 4 x $10^6$ cells/mL in a freshly made solution of imaging media for all experiments. These cell solutions were then allowed to rest at room temperature for 15 min. When the wait time had elapsed, the wash PBS was taken off the well to be imaged, and 200 µL of the neutrophil solution was pipetted into the well. Imaging began as soon as imaging conditions could be verified after placing the cells into the well. All confocal microscopy data for these experiments was collected using the Nikon microscope labeled as “El Capitan”. Care was taken at each magnification to reduce the light exposure used in each experiment, as the swarming process seems more photosensitive than normal migration or calcium behaviors. The camera was run in a 2x2 binning mode. At 10x magnification, all videos are taken with a frame interval of 5s. At 20x and 60x magnification, all videos are taken with either a frame interval of 5 s or 10 s. All videos were taken at 37°C with humidified 5% CO2 for the duration of imaging.

Where indicated, this protocol was modified as follows to add the inhibitors used in this study. For all $cPLA_2$ inhibition experiments, Pyrrophenone was resuspended in DMSO at a stock concentration of 10 mM. This stock was diluted to 500 µM in DMSO, then further to 500 nM (0.5 µM) in imaging media. For the LTB4 inhibitor BIIL315 (Boehringer Ingelheim via opnMe), the drug was resuspended in DMSO at a stock concentration of 10 mM. The stock was diluted to 1 mM in DMSO, then diluted further into imaging media. For the FLAP inhibition experiments, MK886 (MedChemExpress) was diluted from a stock concentration of 10 mM to 1 mM in DMSO, then diluted further into imaging media for a final concentration of 1 µM. Cells were resuspended post pelleting (see above) from culture media and incubated in this drug-media solution for 15 min before starting the experiment. All drugs were kept in solution for the duration of imaging.

When adding exogenous AA to this assay the following modifications were made. First, all vials (Sigma, 4ml amber) used to prepare arachidonic acid solutions were cleaned with a 5% Hellmanex solution, rinsed thoroughly with a stream of milliQ water, and dried using a stream of dry nitrogen. Arachidonic acid supplied in ethanol (Cayman) was first taken from a stock solution at 821 mM and diluted in pure ethanol to 100 mM. This solution was then mixed with imaging media containing 0.5 µM Pyrrophenone to achieve a 10x working concentration of 500 µM of AA and sonicated for at least 10 minutes. Further 10x working dilutions were made using imaging media (+Pyrrophenone) to achieve 50 µM and 10 µM concentrations of AA. Cells were resuspended in 180 uL of Pyrrophenone containing imaging media (no AA), incubated for 15min at RT, then 20 uL of 10x exogenous AA imaging media was added to achieve the final concentrations of 50 µM, 5 µM, and 1 µM of exogenous AA. After AA addition, cells were immediately plated on the swarming targets.

Neutrophil Swarming Accumulation Imaging Protocol:

To measure final accumulation of neutrophils swarming to target sites, the protocol above was modified as followed. Cells were stained and prepared as above, drugs or AA were added as given in the text, and cells were seeded on to targets. Before seeding cells, 60 target positions across 5 wells (12 targets per condition) were found and recorded into a MicroManager multi-position list. Before adding cells, DIC pictures were taken of targets to count the number of yeast at the target site.

Normally, a control, cPLA2 inhibition, cPLA2 inhibition + 1 μM AA, cPLA2 inhibition + 5 μM AA, cPLA2 inhibition + 50 μM AA were all prepared at the same time and run simultaneously. Cells were incubated on the microscope for 1 hour before taking (9 steps, 4 μm per step) z-stack images of the nuclei channel (Hoechst) and center images (single z-plane) for calcium and DIC. All accumulation imaging was done on the Nikon TiE scope labeled "Mt. Diablo" using a Nikon 20 x Plan Apochromat NA 0.75 objective. All experiments were carried out at 37C with humidified 5% CO2 present. The microscope was operated using Micromanager[4].

Light Sheet Nuclei Imaging Protocol:

For light sheet nuclei imaging the neutrophil preparation and swarming assay was modified as follows. After cells were isolated from whole blood (see above), instead of adding both a calcium and Hoechst dye, only a calcium dye was added to the dye step. After cells were spun out of dye media, resuspended in culture media, and plated in a T25; cells were then rested for 1hr. After this wait, SPY-650DNA was added to the culture at 1:1000 dilution from the recommended stock concentration (stock reconstituted to manufacturer specification and aliquoted in single use vials at -20C). Cells were incubated in this culture media + dye mix for 1hr at 37C (with 5% CO2), collected, spun down at 200 x g for 5 min, and resuspended in the same amount of fresh R10 media to achieve a final concentration of 1 x $10^6$ cells/mL.

After this modification of the dye procedure, the swarming assay was setup as normal on the light sheet microscope following the protocol for control samples. minimize thermal, target 8-well chips were incubated in the microscope sample chamber at 37C for at least an hour before cell plating. All images were taken using the Single Objective Light Sheet microscope described below. The microscope was operated using Micromanager[4].

Due to the size and nature of these images, considerable pre-processing was required before cropping, visualization and downstream analysis could take place. Briefly, raw images were run through an algorithm to normalize the data into cartesian space, and large 3D crops were created to enable processing of images with our deconvolution processing pipeline. Full frame ROIs were deconvoluted using 10 iterations of the Richardson-Lucy deconvolution algorithm using an experimentally determined PSF captured though 100nm tetraspeck beads imaged in agarose. These deconvoluted full frame volumes were used for all downstream visualization and analysis.

Arachidonic Acid Micropipette Aspiration Protocol:

The arachidonic acid micropipette assay was arranged as follows. First, 35 mm Mattek dishes and glass vials (4 ml, Amber, Sigma) were washed by hand with a 5% Hellmanex III solution, rinsed with MilliQ water, and then dried under a stream of dry nitrogen. Mattek dishes were then plasma cleaned for 20 minutes and coated with fibronectin immediately following removal from the plasma cleaner. Each dish was coated with 200 uL of imaging media (RPMI + HSA) with 16 ug/ml of porcine fibronectin added, then placed in a humidified incubator at 37C for 30 minutes. Each dish was then thoroughly washed in a stream of MilliQ water, dried with a stream of nitrogen, and left at room temperature until used on the same afternoon of preparation.

Next, a preparation of 20 μM arachidonic acid was prepared fresh each day experiments were carried out. Using cleaned glass vials, an arachidonic acid stock (Cayman) at 821 mM concentration was diluted to 100 mM in pure ethanol. From this working AA stock, a common mixture for control and drug conditions of pipette solution was made by mixing RPMI (without Phenol Red, Gibco) with 3 kDa dextran conjugated to Tetramethylrhodamine (ThermoFisher, final concentration 1 µg/ml) and arachidonic acid working stock to achieve a final concentration of 20 µM. This mixture was then sonicated in a water bath for 15 minutes and separated into different vials for control and drug conditions. To enable drug inclusion in both the cell media and the pipette media, drugs at the concentrations listed for cell incubations were mixed into the pipette arachidonic acid/dye solution. For the ethanol control, pure ethanol was substituted into the pipette solution instead of working arachidonic acid stock and otherwise handled normally.

To load the micropipette with solution, the following steps were followed. First a micromanipulator (Narishige, MM-188NE) was mounted to the Nikon Ti microscope “Mt. Diablo” (detailed description below) using a custom 3d printed mount. 9 µm, polished tip micropipettes (Sunlight Medical Inc) were mounted and secured onto the holder using polystyrene foam. Aspiration was performed manually using the CellTram 4r air system (Eppendorf). Cleaned Mattek dishes (see above, no fibronectin coating) were loaded with 1ml of pipette solution with or without drug depending on condition. The dish was placed on to the scope stage, the micropipette lowered into the solution and liquid was pulled into the pipette via vacuum created by the CellTram. To avoid clogging of the pipette tip, loading took place slightly raised off the bottom of the glass surface, and loading was monitored live via brightfield microscopy. When the pipette had sufficient volume loaded into it, the pipette was raised, and the dish replaced with one with cells on it.

To plate cells on Mattek dishes, neutrophils were spun out of culture (200 x g, 5 min) and resuspended in imaging media with or without drug added. For the drug conditions presented in this work the following final concentrations were used in both cell and pipette solutions: 50 0nM Pyrrophenone, 5 µM MK886, 50 µM Zileuton, 1 µM BIIL315. Cells were resuspended at 1.5 x $10^6$ cells/mL and 200 µL of this solution were placed on to the center of the center glass region of the fibronectin-coated Mattek dish. Careful consideration was taken so that the 200uL drop of cells was centered and only touching the glass part of the dish, and the drop edge pushed with a pipette tip to where the lip of plastic meets the glass coverslip; thus creating an even cell coating in the glass area. Cells were incubated at 37C for 10-15 minutes while the pipette was loaded. Before lowering the pipette into the cell coated dish, 1.5 mL of 37C warmed imaging media was gently pipetted around the inner plastic rim of the dish, forming a donut ring of liquid around the cell drop. After most of the liquid was added, the two liquid interfaces were merged by dragging a pipette tip across the surface until a single liquid interface was achieved; by using this gentle mixing technique, the even cell coating along the glass part of the dish was not disrupted.

Finally, the pipette was lowered into this prepared cell dish, the pipette flow was equilibrated to ensure no media was flowing out of the pipette tip, and then a new position of even density of cells was found. All conditions were imaged on the TiE scope referred to as Mt. Diablo at 37C. 3 channel images (for calcium, nuclei, and pipette dye) were taken at a 5 second interval using a Nikon 20 x Plan Apochromat NA 0.75 objective. Care was taken to always use the same laser and exposure conditions to ensure dextran signal could be roughly comparable between days. Due to the open top nature of these experiments, CO2 exposure was likely limited, however imaging of a single condition never exceeded one hour post cell plating. For each video, the pipette flow was created by turning the CellTram to the same set increase in pressure from the neutral position at 1 minute post recording. Technical replicates were done in the same dish in serial by moving significantly far away from the previous imaging location. A micropipette used in a control condition was sometimes unloaded and reloaded in a following drug condition to cut down on waste, however once a drug solution was used, the pipette was never subsequently reused.

Microscope Equipment Descriptions:

**Nikon Microscope "Mt. Diablo":** Nikon TiE scope configured with a CSU-W1 spinning disk confocal, Borealis beam conditioning unit, Vortran Stradus Versalace laser launch, an air cooled Andor iXon 888 Ultra EM-CCD, TriggerScope 4, and an Okobox temperature/humidity/$CO_2$ controlled environment.

**Nikon Microscope "El Capitan":** Nikon Ti2-E body scope configured with a CrestOptics X-Light V3 confocal spinning disk system, a Lumencor Celesta laser light engine, an Okobox temperature/ humidity/ $CO_2$ controlled environment, Nikon Elements software, and a Photometrics Kinetix sCMOS camera. Where indicated in the text, a Nikon 10x CFI Plan Apo Lambda D objective, Nikon 20x CFI Plan Apo Lambda D NIC N2 objective, or a Nikon 60x Apo TIRF Oil DIC N2 used to achieve the listed magnification. One video in Movie 1 (60x – Yeast Only) was regrettably taken using a Nikon 60x Plan Apo IR WI DIC N2 objective.

**Single Objective Light Sheet Microscope:** This microscope was custom built within the UCSF Center for Advanced Light Microscopy (CALM) following the design principles of Single Objective Light Sheet Microscopy[5,6]. Broadly, this scope included a standard Prior ProScan XYZ stage, Oko stage top incubator (with active humidity control and heated objective collar set), a Prior/Queensgate SP piezo, a VersaLase 8 laser launch, and an ORCA-Quest2 camera. For the optics system, the primary objective (O1) was a CFI SR HP Plan Apo Lambda S 60x Silicon objective; the secondary objective (O2) was a CPI Plan Apochromat Lambda D 40X with a small diameter 170um AR coated coverslip glued to the front (part and service offered by Applied Scientific Instrumentation, SER-GLUE-AR-N40X). The remote 3D image is then collected by a tertiary objective (O3) which is a 'Snouty' lens with a 9mm focal length (AMS-AGY v2) and is tilted away from the optical axis of O2 by 45 and 55 degrees for water and air primary objectives, respectively.

General Image Analysis:

A further discussion of the code used to analyze this data will be provided in the final publication. To request access to the code, please email Evelyn Strickland at, and I will happily supply it to you.

**Method Citations:**

1. Ruoslahti, E., Hayman, E.G., Pierschbacher, M., and Engvall, E. (1982). Fibronectin: purification, immunochemical properties, and biological activities. Methods Enzymol. *82 Pt A*, 803–831.

2. Hopke, A., and Irimia, D. (2020). Ex vivo human neutrophil swarming against live microbial targets. Methods Mol. Biol. *2087*, 107–116.

3. Strickland, E., Pan, D., Godfrey, C., Kim, J.S., Hopke, A., Ji, W., Degrange, M., Villavicencio, B., Mansour, M.K., Zerbe, C.S., et al. (2024). Self-extinguishing relay waves enable homeostatic control of human neutrophil swarming. Dev. Cell *59*, 2659-2671.e4.

4. Edelstein, A.D., Tsuchida, M.A., Amodaj, N., Pinkard, H., Vale, R.D., and Stuurman, N. (2014). Advanced methods of microscope control using Î¼Manager software. J. Biol. Methods *1*, 1.

5. Millett-Sikking, A., and York, A.G. (2019). High NA single-objective lightsheet. Github.io.

6. Sapoznik, E., Chang, B.-J., Huh, J., Ju, R.J., Azarova, E.V., Pohlkamp, T., Welf, E.S., Broadbent, D., Carisey, A.F., Stehbens, S.J., et al. (2020). A versatile oblique plane microscope for large-scale and high-resolution imaging of subcellular dynamics. Elife *9*. https://doi.org/10.7554/eLife.57681.

**Blood Donor Information:**

| Donor #: | Sex: | Age: |
|---|---|---|
| **1** | F | 30 |
| **5** | M | 33 |
| **14** | M | 27 |
| **21** | M | 30 |
| **24** | F | 29 |
| **25** | F | 29 |
| **26** | F | 26 |
| **27** | F | 25 |
| **28** | F | 29 |
| **29** | F | 25 |
| **30** | F | 25 |
| **31** | M | 34 |
| **32** | M | 27 |
| **33** | M | 27 |
